# Arginine valency in *C9ORF72* dipolypeptides mediates promiscuous proteome binding that stalls ribosomes, disable actin cytoskeleton assembly and impairs arginine methylation of endogenous proteins

**DOI:** 10.1101/749127

**Authors:** Mona Radwan, Ching-Seng Ang, Angelique R. Ormsby, Dezerae Cox, James C. Daly, Gavin E. Reid, Danny M. Hatters

## Abstract

*C9ORF72*-associated Motor Neuron Disease patients feature abnormal expression of 5 dipeptide repeat (DPR) polymers. Here we used quantitative proteomics in a Neuro2a cell model to demonstrate that the valency of Arg in the most toxic DPRS, PR and GR, drives promiscuous binding to the proteome, compared to a relative sparse binding of the more inert AP and GA. Notable targets included ribosomal proteins, translation initiation factors and translation elongation factors. PR and GR comprising more than 10 repeats robustly stalled the ribosome suggesting high-valency Arg electrostatically jams the ribosome exit tunnel during synthesis. Poly-GR also bound to arginine methylases and induced hypomethylation of endogenous proteins, with a profound destabilization of the actin cytoskeleton. Our findings point to arginine in GR and PR polymers as multivalent toxins to translation as well as arginine methylation with concomitant downstream effects on widespread biological processes including ribosome biogenesis, mRNA splicing and cytoskeleton assembly.

**SIGNIFICANCE STATEMENT:** The major genetic cause of MND are mutations in an intron of the *C9ORF72* gene that lead to the expansion in the length of a hexanucleotide repeat sequence, and subsequent non-AUG mediated translation of the intron into 5 different DPRs. The two DPRs containing Arg are potently toxic in animal and cell models. Our research shows that the valency of Arg mediates widespread proteome binding especially affecting machinery involved in Arg-methylation, cytoskeleton and translation. We suggest the mechanisms for toxicity are multipronged and involve electrostatic jamming of ribosomes during translation, acting as substrate mimetics for arginine methylase activity that renders the endogenous proteome hypomethylated and impairing actin cytoskeleton assembly. These mechanisms explain pathologic signatures previous reported in human brain pathology.

## INTRODUCTION

The major genetic cause of motor neuron disease (MND) (also known as amyotrophic lateral sclerosis (ALS)) and frontotemporal degeneration (FTD) is an expansion in the number of GGGGCC hexanucleotide repeats in *C9ORF72* from less than 15 in the general population to over 20 (and typically hundreds) (1–4). Toxicity has been proposed to arise through multiple mechanisms including *C9ORF72* haploinsufficiency (2), the formation of *C9ORF72* mRNA foci that sequesters critical RNA binding proteins (5–10) and the production of abnormal translation products by non-AUG-initiated translation (RAN translation) (11). RAN translation occurs from expanded hexanucleotide repeat lengths in both sense and anti-sense transcripts resulting in abnormal expression of 5 dipeptide repeats (DPRs) in neurons of patient brains: poly-GP, poly-GA, poly-GR, poly-AP and poly-PR (5, 8, 12, 13).

Experimental animal and cell culture models expressing the DPRs have revealed poly-GR and poly-PR to be particularly toxic, with the others being comparatively inert (14–17). Furthermore, while all DPRs are widely distributed in human brain of patients with ALS, only poly-GR is correlated to clinically related regions (18). Various interactome studies have indicated that the poly-GR and poly-PR DPRs engage with RNA binding proteins, ribosome machinery and proteins with low complexity domains, which mediate the formation of membrane-less organelles by phase separation (17, 19, 20). These interactions negatively impact on the functioning of ribosome biogenesis (21), ribosome activity (22), nucleolus function (21, 23), nucleocytoplasmic transport (24, 25) and stress granule dynamics (17, 19, 23).

Here we sought to probe the role of the poly-GR and poly-PR DPRs expressed in a simple cell model by defining what they interact with using quantitative proteomics and examining how the valency of short DPR lengths (10× repeats) differed to longer lengths (101× repeats). Our proteomics data suggested potent engagement of poly-PR and poly-GR to ribosome and translational machinery. Given that poly-GR and poly-PR suppress protein translation, we sought to explore the role of the interactions of ribosomes further (17, 19, 20, 22). Here we show that the mechanism to account for disruption of protein translation is by poly-PR and poly-GR with high Arg-valency selectively stalling ribosomes during their synthesis. We also reveal secondary key mechanisms mediating poly-PR and poly-GR toxicity. This includes destabilization of the actin cytoskeleton and proteome arginine hypomethylation. Our findings point to multivalent arginine sequences in the DPRs as promiscuous binders to the proteome that in turn enact multiple modes of toxicity.

## RESULTS

Expression constructs of the DPRs were synthesized using mixed codons and an ATG-start codon, designed to minimize the influence of RNA-mediated effects from the repeat sequences. 10× and 101× repeat lengths of the toxic poly-PR and poly-GR, as well as the less toxic polyGA and polyAP, were prepared.

These constructs displayed patterns of localization and toxicity in the mouse neuroblastoma cell line Neuro2a similar to that described previously in other models (26–31) and observed *in vivo* for cases of MND with *C9ORF72* mutation (5, 8, 13, 32, 33). This included a predominately nuclear punctate pattern of localization for PR_10_ and PR_101_ with residual expression in the cytosol (**Fig 1A**). The longer repeat length accentuated the localization into the nuclear foci. GR_10_ also formed nuclear foci and appeared almost identical to PR_10_, however GR_101_ was excluded from the nucleus and had a mostly diffuse cytoplasmic distribution. By contrast, GA_10_, AP_10_ and AP_101_ were slightly enriched in the nucleus without forming distinct puncta. Similarly, GA_101_ was enriched in the nucleus but also formed large cytosolic (and less commonly nuclear) inclusions in many cells. All DPR constructs expressed at levels comparable to that of GFP except for PR_101_ and GR_101_ that have very weak expression levels; about 30% lower than that of either GFP or other long DPRs (**Fig 1B**). According to cell survival rates when expressing the DPRs, the poly-PR and poly-GR constructs were most toxic and this was true in both 10× and 101× lengths (**Fig 1C**). In contrast, poly-GA was only toxic in 101× lengths and the other DPRS were not toxic.

**Fig 1.**
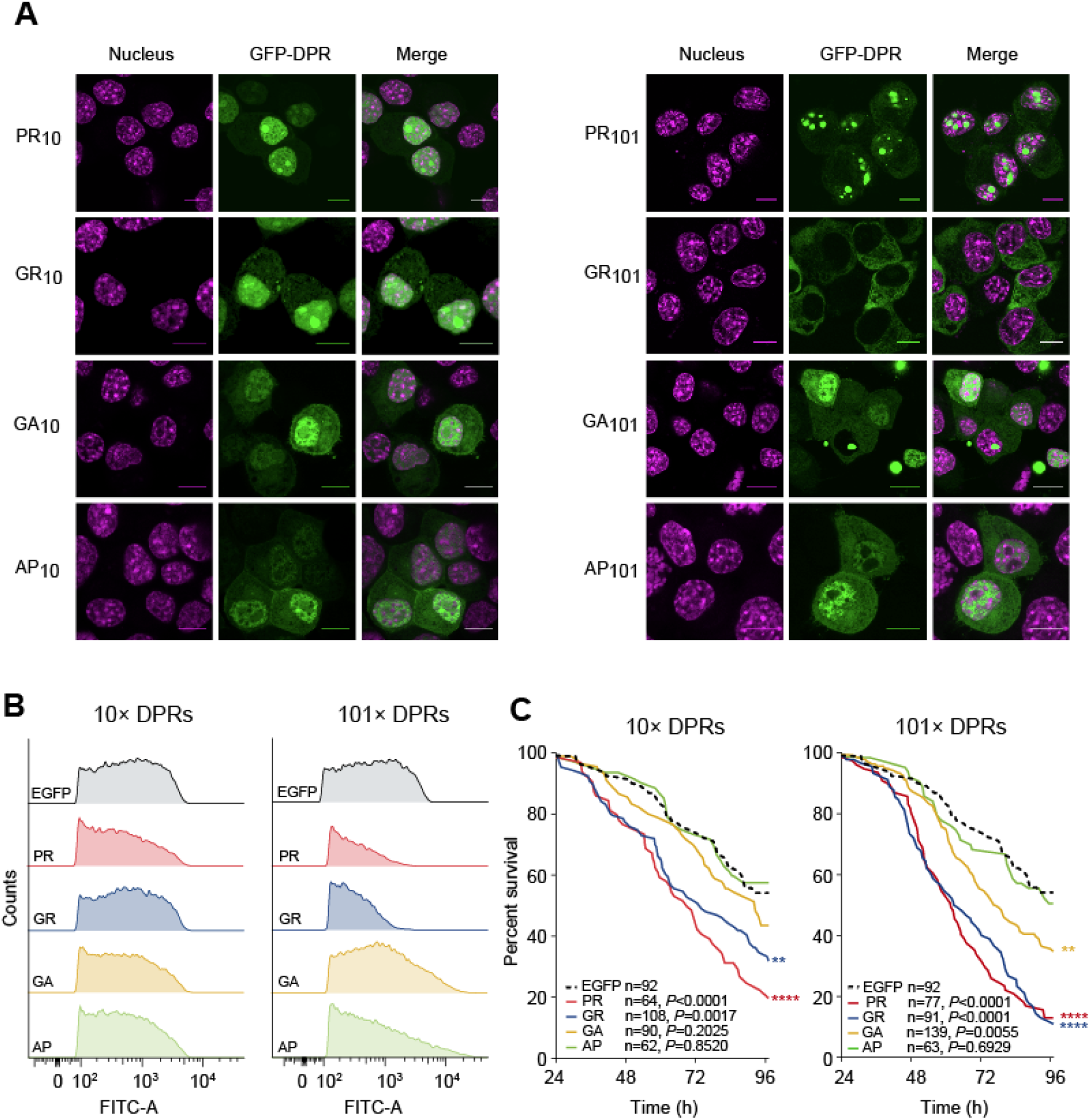
DPR toxicity and cellular localization. **A.** Confocal micrographs of Neuro2a cells 48 h after transfection with GFP-tagged DPRs. The nucleus was stained with Hoechst 33258. Scale bars represent 10 μm. **B.** Expression level of the DPRs as assessed by flow cytometry, 48 h following transfection. FITC-A channel tracks the GFP fluorescence. **C**. Kaplan-Meier survival analysis of Neuro-2a transfected with the GFP-tagged DPRs. Cells were tracked from 24 h post-transfection. *P*-values correspond to log-rank (Mantel-Cox) test of each sample vs EGFP control.

With the model system established, we next sought to investigate how the toxic 101× length DPRs (poly-GR, poly-PR and poly-GA) interacted with the proteome. To do this we transiently transfected cells with GFP or the 101× DPRs fused to GFP and then captured proteins that bound to the DPRs by immunoprecipitation with GFP trap. Proteins were assessed by quantitative proteomics by comparing the pulldowns to GFP-only transfected cells, with protein levels normalized to protein mass recovered from the immunoprecipitation. Under these conditions, GFP levels were anticipated to differ in the pulldown due to different expression levels (**Fig 1B**). Indeed, the amount of GFP appeared heavily enriched in the GFP-only control for the toxic DPRs (PR_101_, GR_101_ and to a lesser extent GA_101_) (**Fig 2A**). Yet, despite the larger amount of GFP coming from the control GFP-only transfected cells (which would enrich for non-specific interactors to GFP), we observed many proteins strongly enriched to the Arg-rich DPRs and comparatively few for GA_101_ and AP_101_ (**Fig 2A**; **Table S1**). The result suggested two conclusions. One was that the Arg appears to mediate promiscuous binding to the proteome and the second was that these interactions are responsible for mitigating toxicity. The Arg-rich DPRs in particular enriched for proteins involved in ribosome biogenesis and RNA splicing machinery, which is consistent with prior findings (17, 34–36). However, we also found additional novel interactions with proteins involved in ribosome-translation, cytoskeleton and chromatin machineries (**Fig 2B**). GR_101_ also enriched specifically for methylosome proteins and PR_101_ with mitochondrial proteins. GA_101_, which is the only DPR that formed large cytosolic inclusions, was enriched for a very distinct proteome, indicative of a quite distinct set of molecular mechanisms involved in its more modest toxicity.

**Fig 2.**
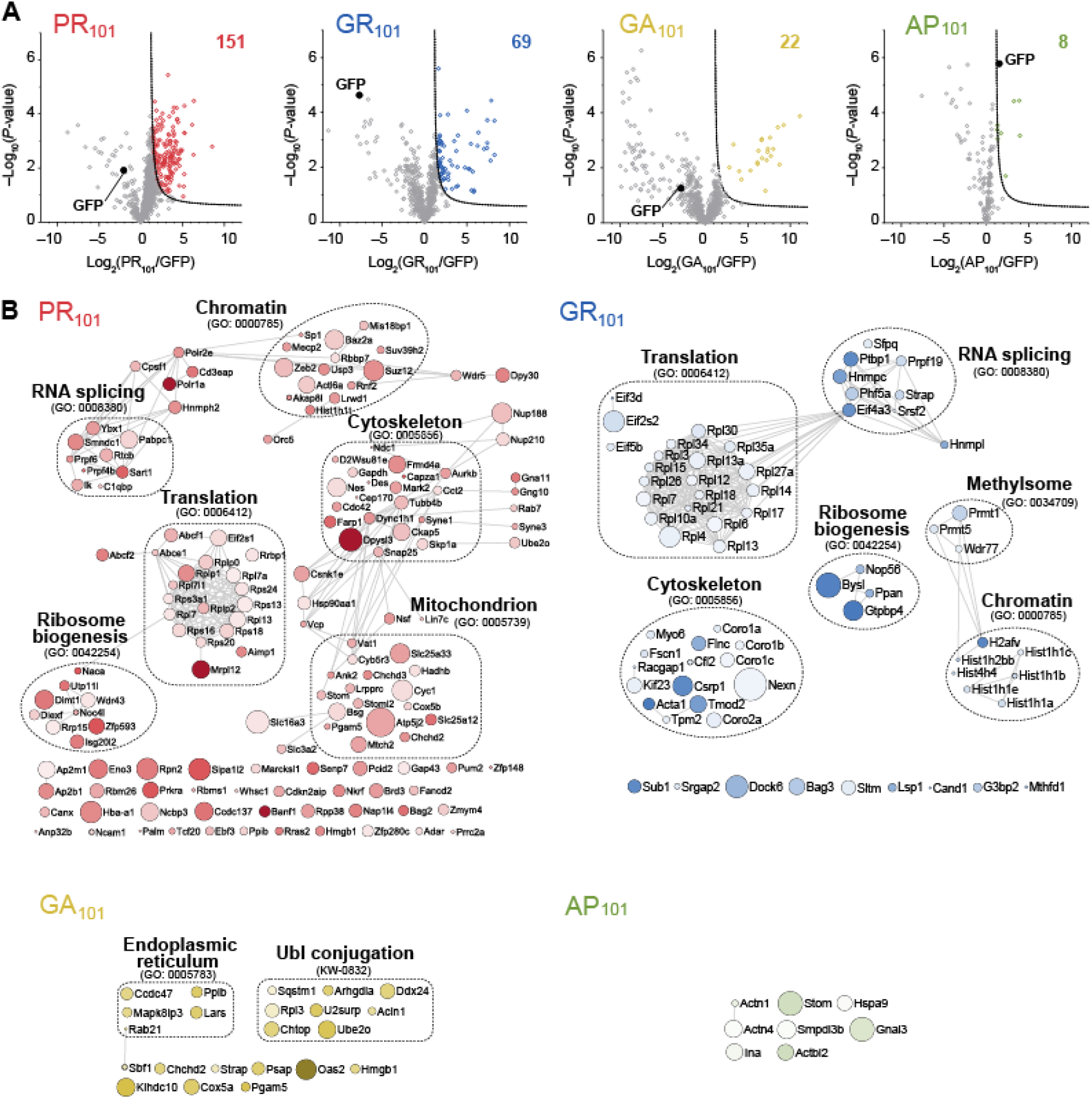
Interactome analysis of the DPR_101_ variants. **A.** Volcano plots of each DPR_101_-GFP versus GFP-only control from quantitative proteomic analysis of GFP-Trap immunoprecipitates of DPR_101_-GFP transfected in Neuro2a cells harvested 48h after transfection. Significant binders (shown in colored circles) were classified with False Discovery Rate of ≤ 0.01 (dotted lines). The number of interactors are indicated. **B.** STRING (v10) interaction maps for proteins significantly enriched with confidence set at 0.9 (highest stringency). Circle sizes are proportional to −log_10_ (*P*-value). The color intensity is proportion to the log (fold change). Selected significantly enriched GO terms (GOCC, GOPB, and UniProt keywords) are displayed.

It was previously reported that Arg-rich DPRs can cause translational suppression although a mechanism that remains undetermined (37). Our proteomics data revealed the ATP-binding cassette sub-family E member 1 (ABCE1) was enriched in the PR_101_ interactome. ABCE1 is involved in translation termination ribosomal recycling (38), which led us to wonder whether synthesis of the Arg-rich DPRs impairs translation. To examine this possibility we employed a previously established assay to measure the ability of a protein sequence to stall or delay protein synthesis rates (39). This assay involves a cassette containing two fluorescent reporters on each side of the peptide sequence to be tested for stalling (GFP at the N-terminus and mCherry at the C-terminus) (**Fig 3A**). Each construct is encoded in frame without stop codons. However the test sequence is flanked by viral P2A sequences, which causes the ribosome to skip the formation of a peptide bond but otherwise continue translation elongation uninterrupted. This means that complete translation of the cassette from one ribosome will generate three independent proteins (GFP, test protein, and mCherry) in an equal stoichiometry. However, should the ribosome stall during synthesis, mCherry is produced at lower stoichiometries than the GFP (**Fig 3A**).

**Fig 3.**
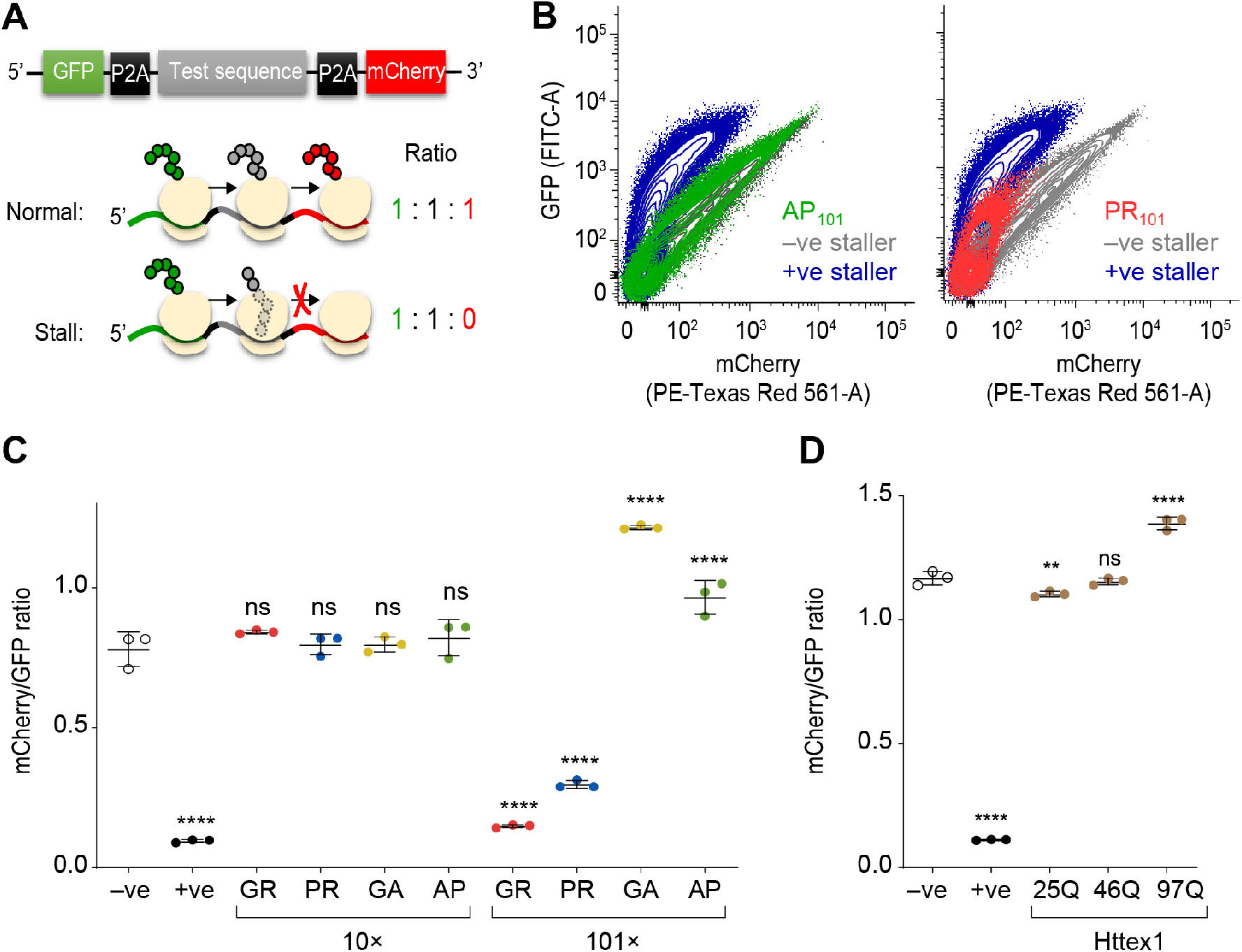
Long Arg-rich DPRs stall ribosomes during translation. **A.** Schematic of the reporter construct design. The P2A sequence causes the ribosome to skip the formation of a peptide bond but otherwise continue translation elongation uninterrupted. Complete translation of the cassette from one ribosome will generate three independent proteins (GFP, test protein, and mCherry). However, should the ribosome stall during synthesis (such as through the previously established positive stall reporter sequence with poly-lysine (K20; +ve sequence. The −ve sequence is stall reporter sequence without lysine residues. **B.** Flow cytograms of Neuro2a cells 48 h after transfection with the indicated test sequences inserted into the reporter. **C.** Median mCherry:GFP ratios calculated from transfected cells population (from 100,000 analyzed cells). Error bars indicate standard deviations from three independent transfections and flow cytometry measurements. *P* values determined for one-way ANOVA and Dunnett’s post hoc test using the -ve as the control. ****, *P*<0.0001; ***, *P*<0.001; ns, *P*>0.05. **D.** The same assay (and statistical tests) using Huntington exon 1 (Httex1) transfected in HEK293 cells with the indicated polyglutamine (polyQ) lengths.

Compared to previously validated FLAG-tagged stalling reporters that either contain 21 AAA codons (which therefore encodes poly-lysine) to stall, or no AAA codons to allow read-through (39), GR_101_ and PR_101_ constructs induced marked stalling (**Fig 3B**). This was DPR-length-dependent in that the 10× repeats showed no stalling (**Fig 3C**). In addition, the AP_101_ and GA_101_ did not lead to stalling (**Fig 3C**). Indeed, there was a significantly increased ratio of mCherry to GFP (**Fig 3C**). A likely explanation for this increased ratio comes from intermolecular FRET arising from low rates of translational readthrough of the stall construct and self-association of these read-through protein products. This was more evident in another control construct of mutant Huntington exon 1 (Httex1), which when containing a polyglutamine (polyQ) expansion above 36 glutamines causes Huntington Disease and becomes highly aggregation prone (40, 41). The expanded polyQ forms of Httex1 did not cause stalling (**Fig 3D**). However, increased pathological lengths of polyglutamine increased the ratio of GFP to mCherry. Microscopic images confirmed the presence of GFP and mCherry in aggregates in cells expressing the GA_101_ and Httex1 reporters (**Fig S1**).

We next sought to examine whether the interactomes of the Arg-rich DPRs were dependent on the valency of arginine (**Fig 4A**). For this analysis, we measured the relative enrichment of the interacting proteins (101× versus 10×) in each GO term significantly associated with the Arg-rich DPRs of both lengths. (The enrichment data of 10× DPRs compared to GFP-control are shown in **Fig S2)**. Almost all GO terms were significantly enriched to the longer Arg-rich DPRs and all were enriched once the relative abundance of the DPRs in the immunoprecipitations was considered (**Fig 4A**). These data therefore suggest arginine valency generally mediates the binding. We also saw enrichment patterns consistent with different cellular localizations of 10× and 101× PR constructs (as shown in **Fig 1**). In particular, the PR_101_ DPR revealed a substantial enrichment of proteins in GO terms relevant to nucleus localization (including DNA replication, positive regulation of transcription) which was consistent with the enhanced localization of PR_101_ to nucleolar substructures compared to PR_10_ (**Fig 4A**). While the actin-related GO terms were enriched also for the Arg-rich DPRs, the enrichment correlated with the DPR localization in the cytosol where most actin is expected to reside. Namely, there was a lesser enrichment for PR_101_ compared to PR_10_, in accordance with the shift from more diffuse nuclear and cytoplasmic localization into the nucleolar substructures. GR_101_ was also excluded from the nucleus compared to GR_10_, and this was reflected in the greater enrichment of GR_101_ with actin related GO terms.

**Fig 4.**
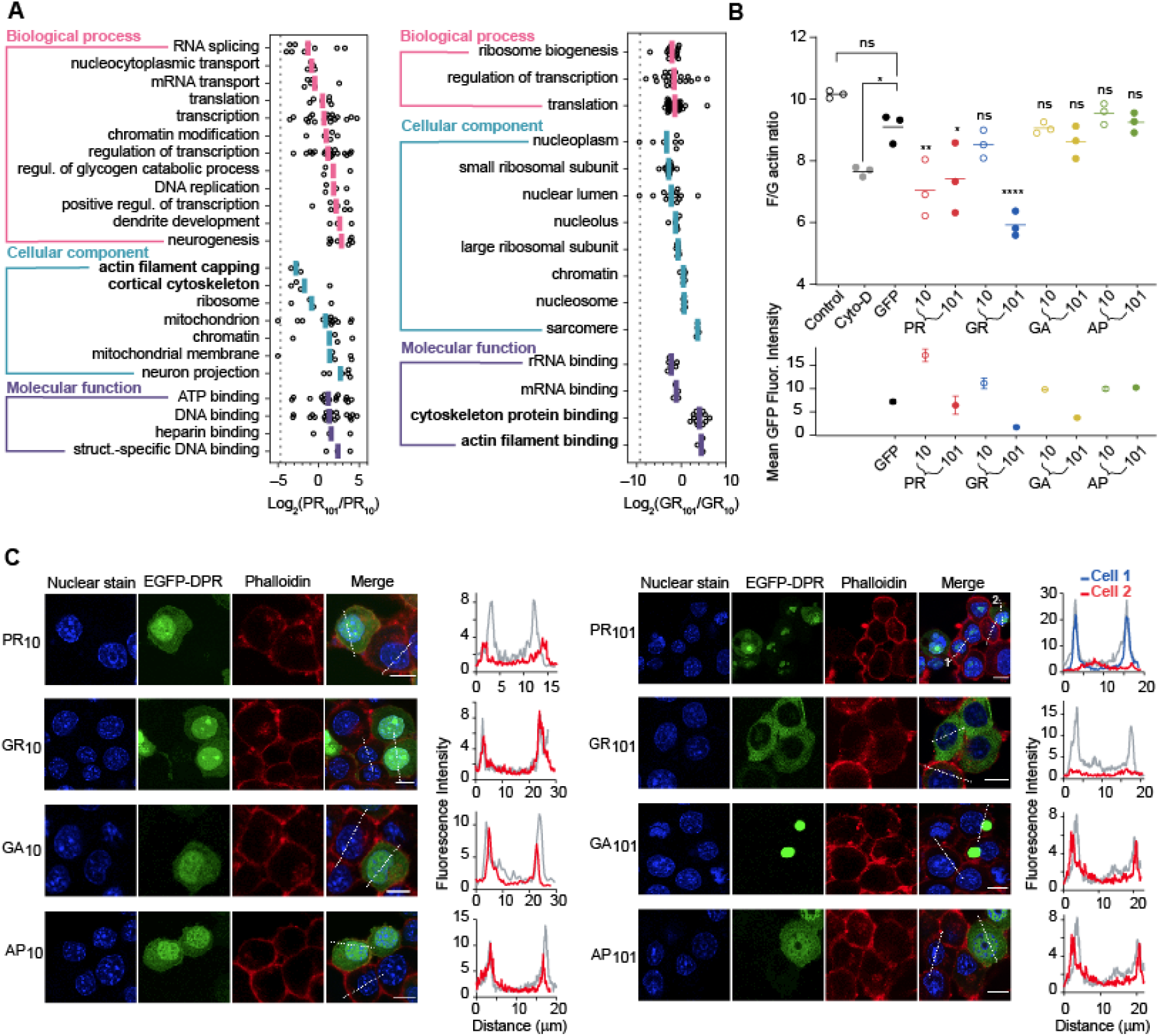
Arg-valency of the DPRs and subcellular location mediate avidity to, and disablement of, actin cytoskeleton assembly. **A.** Gene Ontology (GO) enrichment analysis of proteins significantly associated with the long and short Arg-rich DPRs in the immunoprecipitation. GO terms are shown that contain at least three proteins, and where the log_2_ ratios of DPR length-enrichment values (DPR_101_/DPR_10_), shown as open circles, were deemed different to zero by a Student T-test and Benjamini-Hochberg-adjusted *P* < 0.05. The coloured lines indicate the means of the ratios. The dashed line represents the abundance ratio of GFP, which therefore provides an estimate of the bias of each DPR in the immunoprecipitation. GO terms related to cytoskeleton are shown in bold. **B.** Flow cytometry analyses of population ratios (1×10^4^ Neuro2a cells) of F-actin stain (stained with Phalloidin) and G-actin stain (stained with DNase I) 48 h after transfection with the GFP-tagged DPRs. Cyto-D cells represent 1 h Cytochalasin-D treatment in untransfected cells; Control indicates vehicle control treatment (0.1% DMSO) for Cyto-D sample. The lower graph shows the corresponding mean GFP fluorescence intensities of the samples in the upper graph. Data represent mean ± SD, *N*=3 with *P*-values determined from a one-way ANOVA and Bonferroni’s multiple comparisons post hoc-test with the GFP-alone sample as the control: *, *P*<0.05; **, *P* <0. 01; ****, *P* <0.0001; ns, *P* > 0.05. **C.** Intensity distance trajectories of F-actin (stained with Phalloidin) in Neuro2a cells transfected with GFP-tagged DPRs and nuclei stained with Hoechst 33258. Trajectories are shown on the cells as dotted white lines. Comparisons are shown in the graphs of representative cells (from over 50 measured cells) expressing GFP-tagged DPRs (red or blue lines) and untransfected cells in adjacent cells (grey lines). Scale Bars, 10 μm.

Next, we examined how proteome abundances changed in cells expressing the 100× DPRs. Because the DPRs were toxic and had variable expression levels, we sorted transfected cells into similar levels of expression by virtue of GFP levels before analysis. Cell lysates were collected and quantitatively analyzed by a reductive dimethyl labeling proteome analysis approach (**Fig S3A**). The poly-GR and poly-PR DPRs resulted in the most changes to the proteome in accordance with them having a more potent toxicity and promiscuous pattern of interactions. Poly-GR and poly-PR also had many overlapping GO terms annotated consistent with the Arg-valency driving a similar pathological consequence (**Fig S3B**). Conversely the non-toxic poly-AP resulted in few changes to the proteome abundance and poly-GA had a distinct signature consistent with a different mechanism of toxicity.

Next we examined whether the enrichment with actin GO terms had a functional consequence for the actin cytoskeleton. The Arg-rich DPRs had a significant impact on the formation of filamentous (F) actin compared to the other DPRs and GFP-alone control using a flow cytometry protocol for measuring free and globular (G) actin ratios (**Fig 4B**). There was no apparent co-localization of actin to the DPRs, certainly not to punctate structures of the DPRs (**Fig 4C**). However, it was clear that F actin was reduced in individual cells expressing the Arg-rich DPRs (**Fig 4C**). Therefore, the results collectively suggested that the Arg-rich DPRs led to the binding and destabilization of machinery involved in actin filament assembly.

Arginine methylation has been reported to be abnormal in patients with *C9ORF72* mutations, including the presence of arginine-dimethylated enriched inclusions (42, 43). Arginine residues are commonly methylated to regulate biological activity of many cellular processes and are important in particular in histones which are enriched GO terms in our datasets. In addition, abnormal histone methylation has previously been reported in a mouse model of PR_50_ (44). Furthermore, prior work has suggested that arginine methylase PRMT5 is important in regulating stress granule function in *C9ORF72* models of disease, and methylates ALS-gene risk factor FUS (42). Here, GR_101_ was found to be significantly enriched for 3 proteins in the Methylosome GO term (GO:0034709) including arginine methylases PRMT1 and PRMT5 (**Fig 2B**). PRMT1 appears particularly important, accounting for 85% of the methylation activity in mammalian cells (45).

This led us to hypothesize that the Arg-rich DPRs interfere with endogenous arginine methylation activity, that links to the mechanisms of ALS toxicity. To test this hypothesis, we examined the 101× DPRs affected proteome levels and corresponding levels of arginine methylation. Because of our reductive dimethylation proteomics workflow, we could only observe a minor fraction of possible arginine methylation patterns that could exist. However, sufficient information was obtained to reveal that GR_101_ leads to a significantly lower level of arginine methylation relative to the GFP only control (**Fig 5A**; **Table S3**). These same proteins did not show a significant change in abundance (**Fig 5A**). Examination of these peptides revealed that many come from Hnrnp family proteins including Hnrnpa1, Hnrnpab and Hnrnph1 (**Table S3**). Hnrnpa1 showed a statistically significant reduction in Arg-methylation in the PR_101_ treatment (**Fig 5B**). Hnrnp family proteins are well known substrates of PRMT family proteins (46). Mutations in Hnrnpa1 also cause ALS (47).

**Fig 5:**
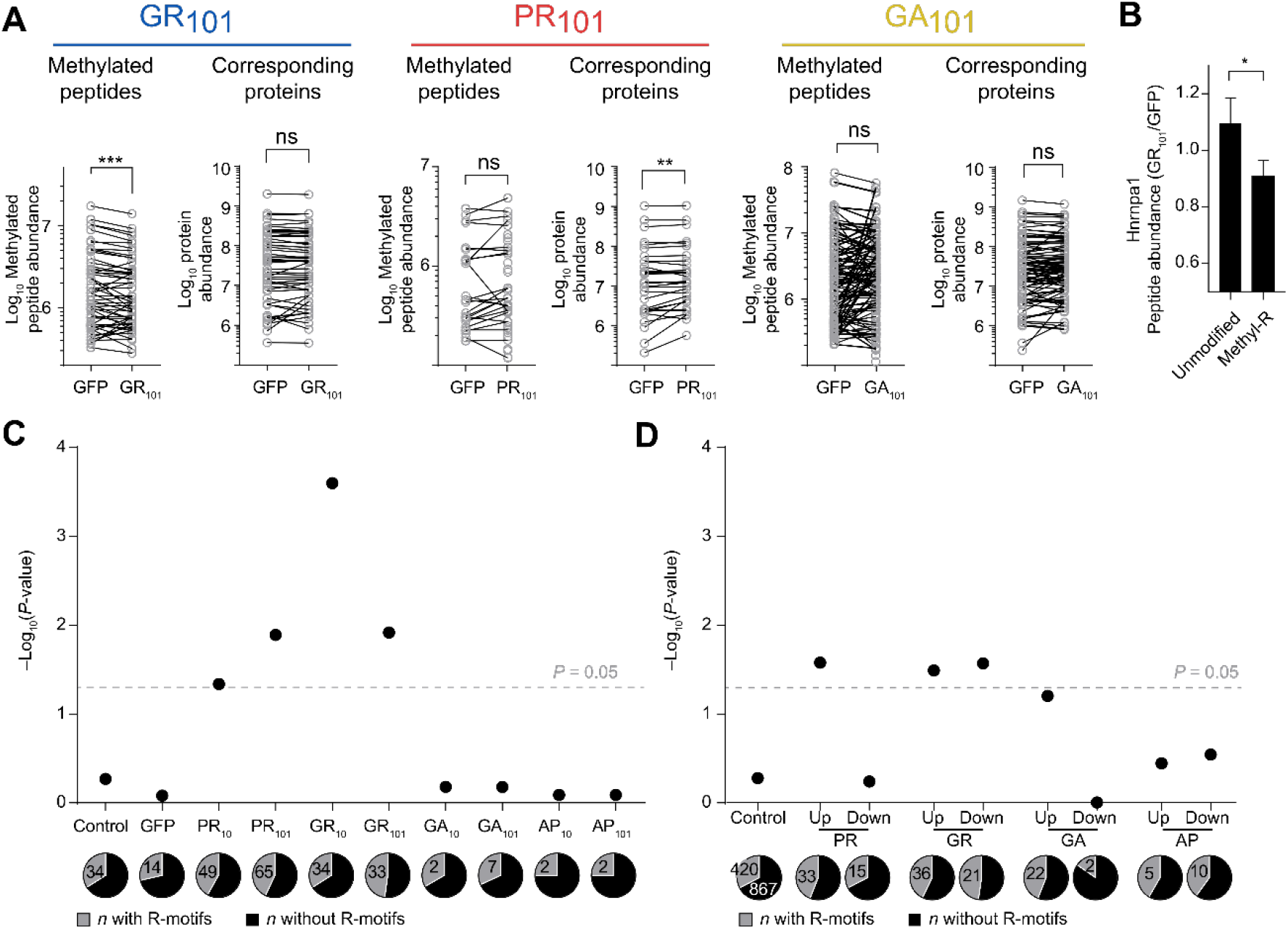
Proteome hypomethylation upon GFP-GR_101_ expression. **A**. Abundance of methylated peptides seen in the whole proteome, and the corresponding protein abundances from which they derive. The data is plotted as matched-pairs of peptides (or proteins) with differences evaluated by 2-tailed Wilcoxon signed rank test. *P* values are coded as ns > 0.05; **, *P* < 0.01; ***, *P* <0.001. The mean difference in abundances of matched pairs of methylated peptides (GR_101_ – GFP) is –399,018. **B.** Shown are the means ± SEM of peptide abundance ratios of Hnrnpa1 from the PR101 and GFP samples. Shown are unmodified and arg-methylated peptides (see **Table S3** for data). Significance of difference was assessed with an unpaired t-test with Welch’s correction. *P* value is coded as *, *P* < 0.05. **C.** Shown are *P* values of a binomial statistical test for DPR interactomes classified into those with arginine-rich motifs (68) or those lacking such motifs. The *P* values indicate the probability of the proportion of proteins with R-motifs being equal to or greater than that observed (versus the control proportion). The control proteins are a dataset of random mouse proteins. **D**. As for C, but for proteins seen significantly upregulated and downregulated in total protein lysate compared to control.

Analysis of the interactome data showed that the Arg-rich DPRs were significantly enriched in interacting with other arginine-enriched proteins (**Fig 5C**). These data suggest the possibility that the Arg-rich DPRs act as substrate sinks of arginine methylases that therefore results in a broader deficiency in arginine methylase modification of endogenous proteins.

## DISCUSSION

Here we show that the Arg-rich DPRs lead to widespread proteome interactions relative to the other less toxic DPRs. These interactions centre on various hubs of cellular activity including translation, ribosome biogenesis, chromatin, mitochondria, cytoskeleton, RNA splicing and the methylsome. We show that the effects appear driven by the valency of arginine as well as changes in the cellular localization of the different lengths of DPRs. In contrast the inert and non-toxic AP DPRs showed few interactions. GA also showed relative few interactors and unmasked a distinct interactome, consistent with a different mechanism of toxicity to the potently toxic Arg-rich DPRs.

Our key finding was that the Arg-rich DPRs manifest toxicity through multiple mechanisms. One mechanism of toxicity is their synthesis leading to a stall in translation. Our study suggests that the Arg-rich DPRs longer than at least 10 repeats are needed to induce stalling. Given that the ribosome exit tunnel holds around 33 amino acids and is lined in negative charges, a plausible explanation for stalling is that DPR lengths approaching 16–17 repeat lengths would be sufficient to fill this cavity volume and lock the peptide in place by electrostatic interactions. Previously it was suggested that a canonical polyadenylate tail on mRNA used to stall ribosomes is translated to lysine, and that the poly-lysine sequence is recognized as aberrant by ribosomes and results in translation repression (48). Additional experiments using repeating sequences of Lys and Arg in proteins both slowed translation, which supports the mechanism of electrostatic interactions jamming the emergent poly-basic chain in the negatively charged ribosome exit tunnel (49). Interestingly, antimicrobial peptides enriched for PR-containing motifs have been demonstrated to bind to the ribosomal exit tunnel and inhibit bacterial protein synthesis (50). This raises the possibility that emergent DPRs can also re-enter and plug the exit tunnel through electrostatic interactions. Further support for this additional mechanism comes from *in vitro* translation assays showing that poly-PR and poly-GR peptides formed insoluble complexes with mRNA, restricted the access of translation factors to mRNA, and blocked protein translation (51). This study showed that poly-PR and poly-GR inhibit protein translation by binding to the translational complex and ribosomal proteins, leading to neurotoxicity (51).

Our data also suggested that the Arg-rich DPRs impede assembly of the actin cytoskeleton. Recently it was observed that promoting actin filament assembly in cell models of ALS can alleviate defects in nuclear-cytoplasmic transport defects (52) which supports the conclusion that destabilization of the actin cytoskeleton is pertinent to disease pathomechanisms. We observed a significant reduction in F-actin in cells with Arg-rich DPRs. Whether this effect is a consequence of other cellular effects, such as hypomethylation or ribosome stalling is unclear. Interestingly it has been reported that arginine methylation of an arginine methylase (PRMT2) regulates the activity of actin nucleator protein Cobl, which suggests a possible role for arginine methylation defects being an upstream mediator of effects on the actin cytoskeleton (53). In addition, it is thought that dysregulation of actin is a key process in ALS (54). Of note is that mutations in profilin-1 protein, which mediates conversion of G-actin to F-actin, is linked to ALS (55). Other cytoskeletal genes have also been linked to ALS including *TUBA4A* and *DCTN1* and *KIF5A* (56, 57).

Our other major finding was that the Arg-rich DPRs led to a broader hypomethylation of the proteome. Previous studies have shown that PRMT1 colocalizes with GR and PR in a *Drosophila* model and that knockdown of PRMT family members enhanced toxicity (58). It was also found that *C9ORF72*-related brain samples had abundant methylated inclusions (58). Thus, the data suggests that the Arg-rich motifs attract and alter the endogenous methylation activity leading to pathological outcomes. The substrate of PRMT family proteins contain glycine- and arginine-rich (GAR) sequences that include multiple arginines in RGG or RXR contexts, which bear resemblance to the Arg-rich DPRs (59). It follows that many of the key pathways seen in our dataset are affected by altered arginine methylase activity – including proteins that are methylated for functional regulation such as histones, proteins involved in mRNA splicing, and ribosomes (60, 61).

Also of note is that other genes that when mutated are risk factors for ALS have activity regulated by arginine methylation and show abnormal methylation patterns in disease. In particular FUS has been reported to interact with PRMT1 and PRMT8 and undergo asymmetric dimethylation in cultured cells (62). Importantly, PRMT1 and PRMT8 localized to mutant FUS-positive inclusion bodies in ALS (62). It has also been reported that arginine methylation modulates the nuclear import of FUS and inclusions in ALS-FUS patients contain methylated FUS (63). We observed the hypomethylation of Hnrnpa1 caused by the arg-rich DPRs, which indicates a possible link to ALS arising from distinct gene mutations. Hnrnpa1 as well as the Arg-rich DPRs and other ALS-associated proteins are known to form molecular condensates by phase separation (57, 64). Arginine methylation of Hnrnpa1 reduces its ability to phase separate suggesting that an imbalance in molecular condensate mechanisms contributes to the pathogenic response (65). PR DPRs can also promote the aggregation of ALS-related proteins containing prion-like domains, that are involved in mediating phase separation into molecular condensates (64). Hence our data indicates a possible convergence of multipronged mechanisms involving methylation, phase separation and cytoskeleton as important contributors to the toxicity of the Arg-rich DPRs.

## AUTHOR CONTRIBUTIONS

M.R. designed and performed the experiments, analyzed the data and helped write the manuscript. C.-S.A. helped perform the proteomics experiments. A.R.O., J.C.D. and D.C. designed and/or performed some of the experiments. G.E.R. oversaw the proteomics experimental design and analysis. D.M.H. oversaw the project, the design of the experiments, interpretation of the data and drafting of the manuscript.

## MATERIALS AND METHODS

An expanded description of the methods are provided in **Supplementary Methods**.

### Plasmids

Synthetic genes for short (10×) and long (101×) dipeptide repeats were synthesized by GeneArt (Life Technologies, Regensburg, Germany) and cloned into pEGFP-C2 to create GFP fusions. The full sequence information is available (**Table S4**).

### Cell culture

Neuro-2a and HEK293T cells, obtained originally from the American Type Culture Collection (ATCC), were maintained in Opti-MEM (Life Technologies) and Dulbecco’s modified Eagle medium (DMEM) (Life Technologies), respectively.

### Flow cytometry

For analysis of DPR expression levels, cells were harvested 48 hr post-transfection and analysed using LSRFortessa X-20 flow cytometer (BD Biosciences). For GFP, data were collected with the 488-nm laser and FITC (530/30) filter. For SYTOX Red dead cell stain, fluorescence was collected using 640-nm laser and APC (670/14) filter.

### Confocal Imaging

Cells expressing GFP-tagged DPRs were fixed 48h after transfection in 4 % paraformaldehyde for 15 min at room temperature. Nuclei were counterstained with Hoechst 33342. Cells were imaged on a Leica SP5 confocal microscope using HCX PL APO CS 40× or 63× oil-immersion objective lens (NA 1.4) at room temperature.

### Longitudinal live cell imaging

Neuro-2a cells in 12-well plate format were co-transfected with individual GFP-tagged DPRs along with mCherry cloned into a pT-Rex vector (Life Technologies). The media was refreshed 24 hours after transfection and cells were then imaged longitudinally with a JuLI stage live cell imaging system with fluorescent images acquired at 15 min intervals for 96 h (Nanoentek, Seoul, South Korea). Death was recorded as the time points at which mCherry fluorescence was lost.

### Sample preparation for proteome analysis of GFP-immunoprecipitated samples

In total 6 × 10^6^ Neuro-2a cells were seeded into 75 cm^2^ flasks and transfected the following day with either GFP-tagged DPRs or GFP-only constructs (24 μg DNA and 60 μL Lipofectamine 2000) according to the manufacturer’s instructions (Life Technologies). Media was refreshed 24h after transfection and cells harvested at 48 h post-transfection by resuspension in ice-cold lysis buffer (10 mM Tris-HCl, pH 7.4; 150 mM NaCl; 0.5 mM EDTA; 0.5% v/v NP-40; 1 mM PMSF; 10 units/ml DNase I) supplemented with EDTA-free Complete protease inhibitor cocktail (Roche Diagnostic). The cell suspensions were then passed through a 27 Gauge syringe needle 25 times, followed by a 31 Gauge needle 10 times and incubated on ice for 30 min. The resultant lysates were clarified by centrifugation (21,000 *g*; 10 min; 4°C). Proteins were immunoprecipitated from 0.5 mg (total protein) of lysate using GFP-Trap MA beads (ChromoTek GmbH, Germany), eluted with 50% v/v aqueous 2,2,2-Trifluoroethanol (TFE), 25mM TEAB and then analysed by mass spectrometry.

### Sample preparation for whole proteome analysis

Neuro2a cells expressing GFP-tagged 101× DPRs were harvested 48 h post transfection in PBS with a cell scraper and gentle pipetting. Cells were pelleted and resuspended, supplemented with DAPI and fluorescence-activated cell sorted using a FACS ARIA III cell sorter (BD Biosciences). 1,000,000 cells of each population of interest were recovered using a matched median GFP expression level. Cells were pelleted and snap frozen in liquid nitrogen.

Cell pellets were thawed then lysed in 100 μL RIPA lysis buffer. Proteins were collected by acetone precipitation and resuspended in 100 μL 0.1M TEAB for mass spectrometry analysis

### Mass spectrometry analysis

Proteins were subjected to reduction with 10 mM tris(2-carboxyethyl)phosphine (TCEP), pH 8.0, and alkylation with 55 mM iodoacetamide for 45 min, followed by trypsin digestion (0.25 μg, 37 °C, overnight). 10 μg of each proteins were labelled by reductive dimethyl labelling using light formaldehyde (CH_2_O), medium formaldehyde (CD_2_O, 98% D) or heavy formaldehyde (^13^CD_2_O, 99% ^13^C, 98% D) for LC-MS/MS analysis.

Samples (1 μg for whole proteome or 0.135 μg for L peptide) were analysed by liquid chromatography-nano electrospray ionization--tandem mass spectrometry (LC-nESI-MS/MS) using Orbitrap Lumos mass spectrometer (Thermo Scientific, San Jose, CA, USA) fitted with nanoflow reversed-phase-HPLC (Ultimate 3000 RSLC, Dionex).

### Proteomic data analysis

For GFP-immunoprecipitated samples, raw MS data were analysed using Proteome Discoverer (version 2.3.0.81; Thermo Fisher Scientific) with the Mascot search engine (Matrix Science version 2.4.1). Data were filtered against the SwissProt *Mus Musculus* database (version 2016_07; 16794 proteins) combined with common contaminant proteins. GFP sequence (UniProt ID: P42212) was also added to the database. For protein identification, the search was conducted with 20 ppm MS tolerance, and 0.8 Da MS/MS tolerance. The false discovery rate (FDR) maximum was set to 1% at the peptide identification level and 5% at the protein identification level. Proteins were filtered for those identified by at least two peptides, one of which was unique, in all three replicates. The common contaminant, Keratin, was excluded from the dataset. Peptide quantitation was performed in Proteome Discoverer v.2.3.0.81.

For whole proteome data analysis, the conditions were similar to those above, but with the following differences: Raw MS data were analysed using Proteome Discoverer (version 2.2; Thermo Fisher Scientific). For protein identification, the search was conducted with 20 ppm MS tolerance, and 0.6 Da MS/MS tolerance. The maximum number of missed cleavage sites permitted was three. Additional variable modifications for mono- and dimethylation of Arginine were included. Peptide quantitation was performed in Proteome Discoverer v.2.2.

### Dual fluorescence translation stall assay

The dual-fluorescence translation stall reporter plasmids were created as described previously, except we used mCherry as the red fluorescent protein moeity (66). In essence, DPR constructs and Httex1 constructs with different polyQ expansions (25Q, 72Q and 97Q) were cloned to replace the linker region. The fluorescence reporter plasmids were transfected into cells using Lipofectamine 2000 and cells were harvested 2-days afterwards for analysis by flow cytometry (LSRFortessa X-20 flow cytometer, BD Biosciences).

### Flow Cytometry analysis of G- and F-actin

F- and G-actin levels in Neuro2a cells were measured as described (67). Briefly, Neuro2a cells were harvested 48 h following transfection of different GFP-tagged DPRs. Cells dosed with 2 μM cytochalasin-D were used as a positive control. Cells incubated with vehicle (0.1% DMSO) were used as negative control (untreated cells). Cells were fixed with 4% paraformaldehyde, permeabilized and stained with with Alexa Flour 594 deoxyribonuclease1 (DNase1) conjugate (10 μg mL^−1^, Invitrogen Molecular Probes, D12372) for G-actin detection and Alexaflour-405 Phalloidin (1:1000, Invitrogen Molecular Probes, A30104) for F-actin detection. Fluorescence was measured using blue (BV421), green (FITC) and red (PE-Texas Red) channels on a FACSCanto flow cytometer (BD Bioscience). A total of 1×10^4^ cells were analysed per acquisition.

### Confocal Microscopy for F-actin analysis

Cells were grown in 8–well ibidi culture chambers (Sarstedt, Nümbrecht, Germany), transfected, fixed with 4% paraformaldehyde and permeabilized with 0.2% Triton X-100/PBS for 5 min. F-actin was stained with Alexa Fluor 594 phalloidin (1:1000, Invitrogen, A12381) for 30 min and with Hoechst 33342 (1:200, Thermo Fisher Scientific) for nuclei staining for 30 min. Cells were imaged on a Leica SP5 confocal microscope using HCX PL APO CS 40× or 63× oil-immersion objective (NA 1.4) at room temperature.

### Statistical analysis

Statistical parameters are reported in the Figures and corresponding Figure Legends. All statistical analyses were performed with GraphPad Prism v 7.05 (Graphpad Software Inc., San Diego, CA, USA).

## Data Availability

Data are available via ProteomeXchange with identifier PXD015180.

## REFERENCES

1. Majounie E, et al. (2012) Frequency of the C9orf72 hexanucleotide repeat expansion in patients with amyotrophic lateral sclerosis and frontotemporal dementia: a cross-sectional study. The Lancet. Neurology 11(4):323–330.

2. DeJesus-Hernandez M, et al. (2011) Expanded GGGGCC hexanucleotide repeat in noncoding region of C9ORF72 causes chromosome 9p-linked FTD and ALS. Neuron 72(2):245–256.

3. Renton AE, et al. (2011) A hexanucleotide repeat expansion in C9ORF72 is the cause of chromosome 9p21-linked ALS-FTD. Neuron 72(2):257–268.

4. Gómez-Tortosa E, et al. (2013) C9ORF72 hexanucleotide expansions of 20–22 repeats are associated with frontotemporal deterioration. Neurology 80(4):366.

5. Gendron TF, et al. (2013) Antisense transcripts of the expanded C9ORF72 hexanucleotide repeat form nuclear RNA foci and undergo repeat-associated non-ATG translation in c9FTD/ALS. Acta neuropathologica 126(6):829–844.

6. Cooper-Knock J, et al. (2014) Sequestration of multiple RNA recognition motif-containing proteins by C9orf72 repeat expansions. Brain : a journal of neurology 137(Pt 7):2040–2051.

7. Sareen D, et al. (2013) Targeting RNA foci in iPSC-derived motor neurons from ALS patients with a C9ORF72 repeat expansion. Science translational medicine 5(208):208ra149–208ra149.

8. Mori K, et al. (2013) The C9orf72 GGGGCC Repeat Is Translated into Aggregating Dipeptide-Repeat Proteins in FTLD/ALS. Science 339(6125):1335–1338.

9. Donnelly CJ, et al. (2013) RNA toxicity from the ALS/FTD C9ORF72 expansion is mitigated by antisense intervention. Neuron 80(2):415–428.

10. Lee Y-B, et al. (2013) Hexanucleotide repeats in ALS/FTD form length-dependent RNA foci, sequester RNA binding proteins, and are neurotoxic. Cell reports 5(5):1178–1186.

11. Zu T, et al. (2011) Non-ATG-initiated translation directed by microsatellite expansions. Proceedings of the National Academy of Sciences of the United States of America 108(1):260–265.

12. Ash PEA, et al. (2013) Unconventional translation of C9ORF72 GGGGCC expansion generates insoluble polypeptides specific to c9FTD/ALS. Neuron 77(4):639–646.

13. Zu T, et al. (2013) RAN proteins and RNA foci from antisense transcripts in C9ORF72 ALS and frontotemporal dementia. Proceedings of the National Academy of Sciences of the United States of America 110(51):E4968–E4977.

14. Mizielinska S, et al. (2014) C9orf72 repeat expansions cause neurodegeneration in Drosophila through arginine-rich proteins. Science (New York, N.Y.) 345(6201):1192–1194.

15. Wen X, et al. (2014) Antisense proline-arginine RAN dipeptides linked to C9ORF72-ALS/FTD form toxic nuclear aggregates that initiate in vitro and in vivo neuronal death. Neuron 84(6):1213–1225.

16. May S, et al. (2014) C9orf72 FTLD/ALS-associated Gly-Ala dipeptide repeat proteins cause neuronal toxicity and Unc119 sequestration. Acta Neuropathologica 128(4):485–503.

17. Lee K-H, et al. (2016) C9orf72 Dipeptide Repeats Impair the Assembly, Dynamics, and Function of Membrane-Less Organelles. Cell 167(3):774–788.e717.

18. Saberi S, et al. (2018) Sense-encoded poly-GR dipeptide repeat proteins correlate to neurodegeneration and uniquely co-localize with TDP-43 in dendrites of repeat-expanded C9orf72 amyotrophic lateral sclerosis. Acta neuropathologica 135(3):459–474.

19. Zhang Y-J, et al. (2018) Poly(GR) impairs protein translation and stress granule dynamics in C9orf72-associated frontotemporal dementia and amyotrophic lateral sclerosis. Nature Medicine 24(8):1136–1142.

20. Moens TG, et al. (2019) C9orf72 arginine-rich dipeptide proteins interact with ribosomal proteins in vivo to induce a toxic translational arrest that is rescued by eIF1A. Acta Neuropathologica 137(3):487–500.

21. Kwon I, et al. (2014) Poly-dipeptides encoded by the C9orf72 repeats bind nucleoli, impede RNA biogenesis, and kill cells. Science 345(6201):1139–1145.

22. Kanekura K, et al. (2016) Poly-dipeptides encoded by the C9ORF72 repeats block global protein translation. Human Molecular Genetics 25(9):1803–1813.

23. Tao Z, et al. (2015) Nucleolar stress and impaired stress granule formation contribute to C9orf72 RAN translation-induced cytotoxicity. Human Molecular Genetics 24(9):2426–2441.

24. Zhang K, et al. (2015) The C9orf72 repeat expansion disrupts nucleocytoplasmic transport. Nature 525(7567):56–61.

25. Freibaum BD & Taylor JP (2017) The Role of Dipeptide Repeats in C9ORF72-Related ALS-FTD. Frontiers in Molecular Neuroscience 10(35).

26. Suzuki H, Shibagaki Y, Hattori S, & Matsuoka M (2018) The proline-arginine repeat protein linked to C9-ALS/FTD causes neuronal toxicity by inhibiting the DEAD-box RNA helicase-mediated ribosome biogenesis. Cell Death Dis 9(10):975.

27. Hartmann H, et al. (2018) Proteomics and *C9orf72* neuropathology identify ribosomes as poly-GR/PR interactors driving toxicity. Life Science Alliance 1(2).

28. Mizielinska S, et al. (2014) C9orf72 repeat expansions cause neurodegeneration in Drosophila through arginine-rich proteins. Science 345(6201):1192–1194.

29. Wen X, et al. (2014) Antisense proline-arginine RAN dipeptides linked to C9ORF72-ALS/FTD form toxic nuclear aggregates that initiate in vitro and in vivo neuronal death. Neuron 84(6):1213–1225.

30. May S, et al. (2014) C9orf72 FTLD/ALS-associated Gly-Ala dipeptide repeat proteins cause neuronal toxicity and Unc119 sequestration. Acta Neuropathol 128(4):485–503.

31. Lee KH, et al. (2016) C9orf72 Dipeptide Repeats Impair the Assembly, Dynamics, and Function of Membrane-Less Organelles. Cell 167(3):774–788 e717.

32. Mori K, et al. (2013) Bidirectional transcripts of the expanded C9orf72 hexanucleotide repeat are translated into aggregating dipeptide repeat proteins. Acta Neuropathologica 126(6):881–893.

33. Schludi MH, et al. (2015) Distribution of dipeptide repeat proteins in cellular models and C9orf72 mutation cases suggests link to transcriptional silencing. Acta Neuropathologica 130(4):537–555.

34. Boeynaems S, et al. (2017) Phase Separation of C9orf72 Dipeptide Repeats Perturbs Stress Granule Dynamics. Molecular Cell 65(6):1044–1055.e1045.

35. Hartmann H, et al. (2018) Proteomics and C9orf72 neuropathology identify ribosomes as poly-GR/PR interactors driving toxicity. Life Science Alliance 1(2).

36. Lin Y, et al. (2016) Toxic PR Poly-Dipeptides Encoded by the C9orf72 Repeat Expansion Target LC Domain Polymers. Cell 167(3):789–802.e712.

37. Moens TG, et al. (2019) C9orf72 arginine-rich dipeptide proteins interact with ribosomal proteins in vivo to induce a toxic translational arrest that is rescued by eIF1A. Acta Neuropathol 137(3):487–500.

38. Pisarev AV, et al. (2010) The role of ABCE1 in eukaryotic posttermination ribosomal recycling. Mol. Cell 37(2):196–210.

39. Juszkiewicz S & Hegde RS (2017) Initiation of Quality Control during Poly(A) Translation Requires Site-Specific Ribosome Ubiquitination. Mol. Cell 65(4):743–750 e744.

40. Duyao M, et al. (1993) Trinucleotide repeat length instability and age of onset in Huntington's disease. Nat. Genet. 4(4):387–392.

41. Scherzinger E, et al. (1999) Self-assembly of polyglutamine-containing huntingtin fragments into amyloid-like fibrils: implications for Huntington’s disease pathology. Proc. Natl. Acad. Sci. U. S. A. 96(8):4604–4609.

42. Chitiprolu M, et al. (2018) A complex of C9ORF72 and p62 uses arginine methylation to eliminate stress granules by autophagy. Nat Commun 9(1):2794.

43. Sakae N, et al. (2018) Poly-GR dipeptide repeat polymers correlate with neurodegeneration and Clinicopathological subtypes in C9ORF72-related brain disease. Acta Neuropathol Commun 6(1):63.

44. Zhang YJ, et al. (2019) Heterochromatin anomalies and double-stranded RNA accumulation underlie C9orf72 poly(PR) toxicity. Science 363(6428).

45. Tang J, et al. (2000) PRMT1 is the predominant type I protein arginine methyltransferase in mammalian cells. J. Biol. Chem. 275(11):7723–7730.

46. Dreyfuss G, Matunis MJ, Pinol-Roma S, & Burd CG (1993) hnRNP proteins and the biogenesis of mRNA. Annu. Rev. Biochem. 62(1):289–321.

47. Kim HJ, et al. (2013) Mutations in prion-like domains in hnRNPA2B1 and hnRNPA1 cause multisystem proteinopathy and ALS. Nature 495(7442):467–473.

48. Ito-Harashima S, Kuroha K, Tatematsu T, & Inada T (2007) Translation of the poly(A) tail plays crucial roles in nonstop mRNA surveillance via translation repression and protein destabilization by proteasome in yeast. Genes Dev. 21(5):519–524.

49. Lu J & Deutsch C (2008) Electrostatics in the ribosomal tunnel modulate chain elongation rates. J. Mol. Biol. 384(1):73–86.

50. Gagnon MG, et al. (2016) Structures of proline-rich peptides bound to the ribosome reveal a common mechanism of protein synthesis inhibition. Nucleic Acids Res 44(5):2439–2450.

51. Kanekura K, et al. (2016) Poly-dipeptides encoded by the C9ORF72 repeats block global protein translation. Hum. Mol. Genet. 25(9):1803–1813.

52. Giampetruzzi A, et al. (2019) Modulation of actin polymerization affects nucleocytoplasmic transport in multiple forms of amyotrophic lateral sclerosis. Nat Commun 10(1):3827.

53. Hou W, et al. (2018) Arginine Methylation by PRMT2 Controls the Functions of the Actin Nucleator Cobl. Dev Cell 45(2):262–275 e268.

54. Hensel N & Claus P (2018) The Actin Cytoskeleton in SMA and ALS: How Does It Contribute to Motoneuron Degeneration? Neuroscientist 24(1):54–72.

55. Wu CH, et al. (2012) Mutations in the profilin 1 gene cause familial amyotrophic lateral sclerosis. Nature 488(7412):499–503.

56. Nicolas A, et al. (2018) Genome-wide Analyses Identify KIF5A as a Novel ALS Gene. Neuron 97(6):1268–1283 e1266.

57. Taylor JP, Brown RH, Jr., & Cleveland DW (2016) Decoding ALS: from genes to mechanism. Nature 539(7628):197–206.

58. Boeynaems S, et al. (2016) Drosophila screen connects nuclear transport genes to DPR pathology in c9ALS/FTD. Sci Rep 6:20877.

59. Blanc RS & Richard S (2017) Arginine Methylation: The Coming of Age. Mol. Cell 65(1):8–24.

60. Yang Y & Bedford MT (2013) Protein arginine methyltransferases and cancer. Nat Rev Cancer 13(1):37–50.

61. Chang FN, Navickas IJ, Chang CN, & Dancis BM (1976) Methylation of ribosomal proteins in HeLa cells. Arch. Biochem. Biophys. 172(2):627–633.

62. Scaramuzzino C, et al. (2013) Protein arginine methyltransferase 1 and 8 interact with FUS to modify its sub-cellular distribution and toxicity in vitro and in vivo. PLoS One 8(4):e61576.

63. Dormann D, et al. (2012) Arginine methylation next to the PY-NLS modulates Transportin binding and nuclear import of FUS. EMBO J. 31(22):4258–4275.

64. Boeynaems S, et al. (2017) Phase Separation of C9orf72 Dipeptide Repeats Perturbs Stress Granule Dynamics. Mol. Cell 65(6):1044–1055 e1045.

65. Ryan VH, et al. (2018) Mechanistic View of hnRNPA2 Low-Complexity Domain Structure, Interactions, and Phase Separation Altered by Mutation and Arginine Methylation. Mol. Cell 69(3):465–479 e467.

66. Juszkiewicz S & S Hegde R (2016) Initiation of Quality Control during Poly(A) Translation Requires Site-Specific Ribosome Ubiquitination.

67. Grosse R, Copeland JW, Newsome TP, Way M, & Treisman R (2003) A role for VASP in RhoA-Diaphanous signalling to actin dynamics and SRF activity. EMBO J 22(12):3050–3061.

68. Mitrea DM, et al. (2016) Nucleophosmin integrates within the nucleolus via multi-modal interactions with proteins displaying R-rich linear motifs and rRNA. Elife 5.

