## Supplementary Methods, Table S4 and Figures for "Arginine valency in *C9ORF72* dipolypeptides mediates promiscuous proteome binding that stalls ribosomes, disable actin cytoskeleton assembly and impairs arginine methylation of endogenous proteins"

- **Table S1: Protein interactors to 101× DPRs. Relates to Fig 2 & Fig S2.** *Separate file*
- **Table S2: Cellular abundances of proteins caused by 101× DPR expression. Relates to Fig S3.** *Separate file*
- **Table S3: Arginine methylated proteome quantitation by 101× DPR expression. Relates to Fig 5.** *Separate file*
- **Table S4: Sequence of the open reading frames from the synthetic DPR expression constructs.**

- **Fig S1: Expression of DPRs in the ribosome stall reporter constructs.**
- **Fig S2. Interactome analysis of the $DPR_{10}$ variants. Relates to Fig 2.**
- **Fig S3: Proteome abundance changes in response to $DPR_{101}$ expression. Relates to Fig 4.**

- **Supplementary References.**

### Supplementary Methods

#### Plasmids

Synthetic genes for short (10×) and (101×) dipeptide repeats were synthesized by GeneArt (Life Technologies, Regensburg, Germany). The full sequence information is available (**Table S4**). These constructs were flanked 5' by XhoI and PstI recognition sites and 3' by BclI and BamHI recognition sites. XhoI and BamHI enzymes (NEB) were used to introduce the gene cassette into the mammalian expression vector pEGFP-C2 vector so the GFP tag was on the N-terminus of the 10× repeat sequence. The 101× DPR sequence were inserted between PstI and BclI sites in the backbones of the previously developed pEGFP-C2-10× DPR via classical restriction digest cloning following manufacturers' recommendations.

#### Cell culture

Neuro-2a and HEK293T cells, obtained originally from the American Type Culture Collection (ATCC), were maintained in Opti-MEM (Life Technologies) and Dulbecco's modified Eagle medium (DMEM) (Life Technologies), respectively. The medium was supplemented with 10% v/v fetal calf serum, 1 mM glutamine, and 100 Unit $mL^{-1}$ penicillin and 100 µg $mL^{-1}$ streptomycin, and cells were kept in a humidified incubator with 5% v/v atmospheric $CO_2$.

#### Flow cytometry

For analysis of DPR expression levels, cells were harvested 48 hr post-transfection, in 500 µL PBS containing 0.5 µL of 5 µM SYTOX Red dead cell stain (Invitrogen). Cells were analysed using LSRFortessa X-20 flow cytometer (BD Biosciences). A minimum of 100,000 events per sample were collected at a high flow rate. Forward scatter and side scatter were collected. For GFP, data were collected with the 488-nm laser and FITC (530/30) filter. For SYTOX Red dead cell stain, fluorescence was collected using 640-nm laser and APC (670/14) filter. Flow cytometric gating and data analysis was performed using FlowJo software (v10.5.3) and graphs were analysed in GraphPad Prism 7.05.

#### Confocal Imaging

Cells expressing GFP-tagged DPRs were fixed 48 h after transfection in 4 % paraformaldehyde for 15 min at room temperature. Nuclei were counterstained with Hoechst 33342 at 1:200 dilution (Thermo Fisher

Scientific) for 30 min then washed twice in PBS. Fixed cells were imaged on a Leica SP5 confocal microscope using HCX PL APO CS 40× or 63× oil-immersion objective lens (NA 1.4) at room temperature. The Hoechst 33342 channel was collected with an excitation wavelength of 405 nm and emission wavelengths of 445–500 nm; EGFP was collected by excitation at 488 nm and emission from 520–570 nm. Single colour controls were used to establish and adjust to remove bleed through of the emission filter bandwidths. FIJI version of ImageJ (1) and Inkscape were used for image processing.

**Longitudinal live cell imaging**

Neuro-2a cells in 12-well plate format were co-transfected with individual GFP-tagged DPRs along with mCherry in a pT-Rex vector (Life Technologies). The media was refreshed 24 hours after transfection and cells were then imaged longitudinally with a JuLI stage live cell imaging system with fluorescent images acquired at 15 min intervals for 96 h (Nanoentek, Seoul, South Korea). Channels used: GFP for EGFP (Excitation: 466/40, Emission: 525/50), RFP for mCherry (Excitation: 525/50, Emission: 580 LP).
Death was recorded as the time points at which mCherry fluorescence was lost. This event corresponded to the loss of membrane integrity and cell death and was found to be a highly sensitive and specific assay of cell death through different pathways and in different types of cell (2, 3). Cells that drifted from focus were censored. For statistical analysis, survival time was defined as the imaging time point at which a cell was last seen alive. Kaplan–Meier curves were used to estimate survival and hazard functions with GraphPad Prism software. Differences in Kaplan–Meier curves were assessed with Log-rank (Mantel-Cox) test.

**Sample preparation for proteome analysis of GFP-immunoprecipitated samples**

$6 \times 10^6$ Neuro-2a cells were seeded into 75 $cm^2$ flasks and transfected the following day with either GFP-tagged DPRs or GFP-only constructs (24 µg DNA and 60 µL Lipofectamine 2000) according to the manufacturer's instructions (Life Technologies). The experiment was designed as 3 biological replicates. Media was refreshed 24h after transfection. At 48 h post-transfection, cells were gently rinsed with PBS and harvested in PBS by gently pipetting. Cells were pelleted (120 *g*; 6 min; room temperature) and resuspended in 1 mL PBS and pelleted again (400 *g*; 6 min; room temperature). The pellet was resuspended in 200 µL ice-cold lysis buffer (10 mM Tris-HCl, pH 7.4; 150 mM NaCl; 0.5 mM EDTA; 0.5% v/v NP-40; 1 mM PMSF; 10 units/ml DNase I) supplemented with EDTA-free Complete protease inhibitor cocktail (Roche Diagnostic). The cell suspensions were then passed through a 27 Gauge syringe needle

25 times, followed by a 31 Gauge needle 10 times and incubated on ice for 30 min. The resultant lysates were clarified by centrifugation (21000 *xg*; 10 min; 4°C). Protein concentrations were quantified by the Pierce BCA Protein Assay (Catalogue Number: 23225, Thermo Fischer Scientific, MA, USA) using bovine serum albumin (BSA) as the mass standard. 0.5 mg of cellular protein was added to 25 µL of GFP-Trap MA beads (ChromoTek GmbH, Germany) pre-washed and equilibrated in the wash buffer (10 mM Tris/Cl pH 7.4; 150 mM NaCl; 0.5 mM EDTA; 1 mM PMSF; EDTA-free Protease inhibitor cocktail). The solution was incubated for 2 hr at 4°C (end-over-end rotation). Magnetically separated beads were then washed 3 times with wash buffer and 2 times more with 25 mM triethylammonium bicarbonate (TEAB) buffer. The immunoprecipitated proteins were eluted by the addition of 100 µl of 50% v/v aqueous 2,2,2-Trifluoroethanol (TFE), 25mM TEAB. The supernatant was collected after pelleting (2000 *g*; 2 min; room temperature) and adjusted to a final concentration of 100 mM TEAB by addition of 1 M stock solution (and the pH was validated to be approximately 7 after this treatment). The samples were further processed for mass spectrometry analysis.

**Sample preparation for whole proteome analysis**

Neuro2a cells expressing GFP-tagged 101× DPRs were harvested 48 h post transfection in PBS with a cell scraper and gentle pipetting. Cells were pelleted (120 *g*; 6 min) and resuspended in 2 ml PBS supplemented with 10 units $mL^{-1}$ DNase I and filtered through 100-µm nylon mesh before analysis by flow cytometry. Just before sorting, 2 µL of the nuclear marker DAPI (1:1000, D1306, Thermo Fisher Scientific) was spiked into cell suspensions to stain dead cells. Cells were sorted using a FACS ARIA III cell sorter (BD Biosciences) equipped with 405-nm, 488-nm, 561-nm and 640-nm lasers using a 100-µm nozzle. Gating was performed with BD FACS Diva software (Becton, Dickinson Biosciences). Cells (1,000,000) of each population of interest were sorted at a speed of 1500 cells/s. Side scatter (SSC) and forward scatter (FSC) height, width, and area were collected to gate for the single cell population. DAPI area was collected to gate-out dead cells. Data were also collected for pulse height, width, and area of GFP with the FITC filter. To match for expression, cells were further gated to the same median GFP intensity of 2200 fluorescence units by varying the window of expression. Cells were sorted in parallel across three days and performed as three matched replicates. Cells were kept on ice for all steps of the sorting preparation and handling. The targeted population was directly sorted into PBS, pelleted (120 *g*, 6 min), and snap frozen in liquid nitrogen then kept at –80 °C until use.

Cell pellets were thawed and resuspended in 100 µL RIPA lysis buffer (25mM Tris-HCl, pH 7.4, 150mM NaCl, 1% NP40, 0.1% SDS, 1% Sodium deoxycholate, 1× complete mini-protease cocktail; Roche), and incubated on ice for 30 min. The concentration of proteins was measured by the Pierce BCA Protein Assay according to the manufacturer's instruction (Thermo Fisher Scientific). Equal amounts of protein for each sample were precipitated with six volumes of pre-chilled (–20°C) acetone, and incubation overnight. Samples were pelleted at 21,000 *g* at 4°C for 10 min. Acetone was decanted without disturbing the protein pellet. The pellets were washed once with pre-chilled acetone then allowed to dry for 10 min. The protein precipitates were resuspended in 100 µL 0.1M TEAB and were vortexed and sonicated 3 times for 30 sec to help solubilize the pellet. The samples were further processed for mass spectrometry analysis

**Mass spectrometry analysis**

Proteins were subjected to reduction with 10 mM tris(2-carboxyethyl)phosphine (TCEP), pH 8.0, and alkylation with 55 mM iodoacetamide for 45 min, followed by trypsin digestion (0.25 µg, 37 °C, overnight). The resultant peptides were then desalted by solid-phase extraction following acidification in 1% v/v formic acid; the cartridge (Oasis HLB 1 cc Vac Cartridge, product number 186000383, Waters Corp., USA) was pre-washed with 1 mL of 80% v/v acetonitrile (ACN) containing 0.1% v/v trifluoroacetic acid (TFA) and equilibrated with 1.2 mL of 0.1% v/v TFA three times. Samples were then loaded on the cartridge and washed with 1.5 mL of 0.1% v/v TFA before being eluted with 0.8 ml of 80% v/v ACN containing 0.1% v/v TFA and collected in 1.5 mL microcentrifuge tubes. Peptides were then lyophilized by freeze drying (Virtis, SP Scientific). The peptides were resuspended in 100 µL distilled water and quantified using microBCA assay (Catalogue Number: 23235, Thermo Fischer Scientific) with BSA as the mass standard. Then, 10 µg of each sample (in a volume of 50 µL containing 100 mM TEAB) were differentially labelled by reductive dimethyl labelling using equal volumes (2 µL) of 4% light formaldehyde ($CH_2O$), 4% medium formaldehyde ($CD_2O$, 98% D) or heavy formaldehyde ($^{13}CD_2O$, 99% $^{13}C$, 98% D) and 0.6 M Sodium cyanoborohydride ($NaCNBH_3$, for light and medium label) or Sodium cyanoborodeuteride ($NaCNBD_3$, 96% D, for heavy label) added in sequence. The peptide solutions were incubated on an Eppendorf Thermomixer at room temperature for 1 h. After quenching with 8 µL of 1% v/v ammonium hydroxide followed by 8 µL of neat formaldehyde, dimethyl-labelled peptides were mixed up in equal volumes before LC-MS/MS analysis.

Samples were analysed by liquid chromatography-nano electrospray ionization--tandem mass spectrometry (LC-nESI-MS/MS) using Orbitrap Lumos mass spectrometer (Thermo Scientific, San Jose, CA, USA) fitted with nanoflow reversed-phase-HPLC (Ultimate 3000 RSLC, Dionex). The nano-LC system was equipped with an Acclaim Pepmap nano-trap column (Dionex – C18, 100 Å, 75 µm × 2 cm) and an Acclaim Pepmap RSLC analytical column (Dionex – C18, 100 Å, 75 µm × 50 cm). For each LC-MS/MS experiment, 1 µg (whole proteome) or L (0.135 ug peptide) of the peptide mix was loaded onto the enrichment (trap) column at an isocratic flow of 5 µL/min of 3% $CH_3CN$ containing 0.1% formic acid for 6 min before the enrichment column is switched in-line with the analytical column. The eluents used for the LC were 5% DMSO/0.1% v/v formic acid (solvent A) and 100% $CH_3CN$/5% DMSO/0.1% formic acid v/v. The gradient used was 3% B to 20% B for 95 min, 20% B to 40% B in 10 min, 40% B to 80% B in 5 min and maintained at 80% B for the final 5 min before equilibration for 10 min at 3% B prior to the next analysis.

The mass spectrometer was operated in positive-ionization mode with spray voltage set at 1.9 kV and source temperature at 275°C. Lockmass of 401.92272 from DMSO was used. The mass spectrometer was operated in the data-dependent acquisition mode MS spectra scanning from m/z 400–1500 at 120,000 resolution with AGC target of $5e^5$. The "top speed" acquisition method mode (3 sec cycle time) on the most intense precursor was used whereby peptide ions with charge states ≥2-5 were isolated with isolation window of 1.6 m/z and fragmented with high energy collision (HCD) mode with stepped collision energy of 30 ±5%. Fragment ion spectra were acquired in Orbitrap at 15000 resolution. Dynamic exclusion was activated for 30s.

**Proteomic data analysis**

For GFP-immunoprecipitated samples, raw MS data were analysed using Proteome Discoverer (version 2.3.0.81; Thermo Fisher Scientific) with the Mascot search engine (Matrix Science version 2.4.1). Data were filtered against the SwissProt *Mus Musculus* database (version 2016_07; 16794 proteins) combined with common contaminant proteins. GFP sequence (UniProt ID: P42212) was also added to the database. For protein identification, the search was conducted with 20 ppm MS tolerance, and 0.8 Da MS/MS tolerance. The enzyme specificity was set as trypsin. The maximum number of missed cleavage sites permitted was two, and the minimum peptide length required was six. The following modifications were allowed: Oxidation (M), Acetylation (Protein N-term), Dimethylation (K), Dimethylation (N-Term), Dimethylation: 2H(4) (K), Dimethylation 2H(4) (N-term), 2H(6)13C(2)

Dimethylation (K), 2H(6)13C(2) Dimethylation (N-term) (Variable); Carbamidomethyl (C) (Fixed). The false discovery rate (FDR) maximum was set to 1% at the peptide identification level and 5% at the protein identification level. Proteins were filtered for those identified by at least two peptides, one of which was unique, in all three replicates. The common contaminant, Keratin, was excluded from the dataset.

Peptide quantitation was performed in Proteome Discoverer v.2.3.0.81 using the precursor ion quantifier node. Dimethyl labelled peptide pairs (between two comparison of light, medium or heavy) were established with a 2 ppm mass precision, and a signal to noise threshold of 3. A retention time tolerance of isotope pattern multiplets was set to 0.8 min. Three single peak or missing channels were allowed for peptide identification. The protein abundance in each replicate was calculated by summation of the unique peptide abundances that were used for quantitation (light, medium and-or heavy dimethyl derivatives). Missing quantitation values was replaced with a constant (zero-filling). The peptide group abundance and protein abundance values were normalized to account for sample loading. In brief, the total peptide abundances for each sample was calculated and the maximum sum for all files was determined. The normalization factor was the factor of the sum of the sample and the maximum sum in all files. After calculating the normalization factors, the Peptide and Protein Quantifier node normalized peptide group abundances and protein abundances by dividing abundances with the normalization factor over all samples.

The normalized protein abundances were imported into Perseus software (v 1.6.5.0). Protein abundances were transformed to $log_2$ scale. The samples were then grouped according to the replicates. For pairwise comparison of proteomes and determination of significant differences in protein abundances Welch's t-test based on permutation-based FDR statistics was then applied (250 permutations; FDR=0.01; S0=1) were applied.

For whole proteome data analysis, the conditions were similar to those above, but with the following differences: Raw MS data were analysed using Proteome Discoverer (version 2.2; Thermo Fisher Scientific). For protein identification, the search was conducted with 20 ppm MS tolerance, and 0.6 Da MS/MS tolerance. The maximum number of missed cleavage sites permitted was three. Additional variable modifications for mono- and dimethylation of arginine were included. Peptide quantitation was performed in Proteome Discoverer v.2.2, and the retention time tolerance of isotope pattern multiplets was set to 0.6 min. After grouping samples, proteins with at least four valid values across all groups were retained for the subsequent analysis. Missing values were imputed from a distribution of all other $log_2$-transformed protein values from that sample, using the default settings in Perseus (1.8 standard deviation downshift, 0.3 mean down shift). For pairwise comparison of proteomes and determination of significant differences in protein abundances, a two-sample Students t-test based on permutation-based FDR statistics was applied (250 permutations; FDR=0.05; S0=0.1) were applied.

**Dual fluorescence translation stall assay**

Genes were synthesized to produce the dual-fluorescence translation stall reporter as described previously (4), except we used mCherry as the red fluorescent protein. DPR constructs, or Httex1 with different polyQ expansions (25Q, 72Q and 97Q) were cloned to replace the linker region using PstI and BamHI restriction sites. Frame shifts were corrected using standard PCR-based strategies and were validated by sequencing. Dual fluorescence reporter plasmids were transfected into cells using Lipofectamine 2000 (Life Technologies) according to manufacturer guidelines. Two days after transfection, cells were harvested in PBS containing SYTOX Blue dead cell stain (S34857, Thermo Fischer Scientific) to exclude dead cells from subsequent analysis. Cellular fluorescence of 100,000 events per sample were analysed on a LSRFortessa X-20 flow cytometer (BD Biosciences) using the 488-nm laser and 530/30 filter for GFP and the 561nm laser and 610/20 bandpass filter for mCherry. Subsequent analysis of flow cytometry data was done using FlowJo software (v10.5.3) and graphs were analysed in GraphPad Prism 7.05.

**Flow Cytometry analysis of G- and F-actin**

F- and G-actin levels in Neuro2a cells were measured as described (5). Briefly, Neuro2a cells seeded into 12-well plates were harvested 48 h following transfection of different GFP-tagged DPRs. Cells were treated with 2 µM cytochalasin-D in 0.1% DMSO/PBS for 1 h for the positive control. Cells incubated with 0.1% DMSO for 1 h were used as negative control (untreated cells). Cells were fixed with 4% paraformaldehyde for 15 min, then permeabilized with 0.2% Triton X-100 for 5 min. After washing in PBS, cells were blocked with 1% BSA in 0.1% v/v Triton X-100/PBS solution for 15min, and then incubated in the dark at room temperature for 30 min with Alexa Flour 594 deoxyribonuclease1 (DNase1) conjugate (10 µg $mL^{-1}$, Invitrogen Molecular Probes, D12372) for G-actin detection and Alexaflour-405 Phalloidin (1:1000, Invitrogen Molecular Probes, A30104) for F-actin detection. After thorough washing with 1× PBS, fluorescence was measured using blue (BV421), green (FITC) and red (PE-

Texas Red) channels on a FACSCanto flow cytometer (BD Bioscience). A total of $1\times10^4$ cells were analysed per acquisition. Unstained cells were used to set the baseline. G- and F-actin contents were determined from the respective fluorescence and the ratio of F/G calculated from the mean values as determined with FlowJo software.

**Confocal Microscopy for F-actin analysis**

Cells were grown in 8–well ibidi culture chambers (Sarstedt, Nümbrecht, Germany). Cells were washed with PBS once before being fixed with 4% paraformaldehyde for 15 min at room temperature, then washed with PBS 3 times and permeabilized in 0.2% w/v Triton X-100/PBS for 5 min. The cells were then blocked with 1% w/v BSA in 0.1% w/v Triton X-100/PBS for 15 min at room temperature. F-actin was stained with Alexa Fluor 594 phalloidin (1:1000, Invitrogen, A12381) for 30 min and with Hoechst 33342 (1:200, Thermo Fisher Scientific) for nuclei staining for 30 min in the dark. Cells were imaged on a Leica SP5 confocal microscope using HCX PL APO CS 40× or 63× oil-immersion objective (NA 1.4) at room temperature. Fluorescence intensity profiles of phalloidin in >50 cells expressing GFP-tagged DPRs and untransfected cells were analysed using ImageJ software.

**Statistical analysis**

Statistical parameters are reported in the Figures and corresponding Figure Legends. All statistical analyses were performed with GraphPad Prism v 7.05 (Graphpad Software Inc., San Diego, CA, USA).

**Data Availability**

The mass spectrometry proteomics data have been deposited to the ProteomeXchange Consortium via the PRIDE (6) partner repository with the dataset identifier PXD015180.

**Table S4. Sequence of the open reading frames from the synthetic DPR expression constructs.**

| Construct | Sequence |
|---|---|
| **10x DPRs** | |
| $GA_{10}$ | 5’-GGCGCTGGCGCTGGGGCAGGCGCAGGGGCTGGCGCAGGCGCTGGGGCTGGGGCTGGGGCA-3’ |
| $GR_{10}$ | 5’-GGCAGAGGAAGAGGCAGGGGACGCGGAAGGGGGAGAGGACGCGGCAGAGGCCGGGGAAGA-3’ |
| $AP_{10}$ | 5’-GCACCAGCTCCAGCCCCTGCTCCTGCTCCCGCCCCAGCACCCGCCCCTGCCCCAGCCCCA-3’ |
| $PR_{10}$ | 5’-CCCAGACCTAGACCTCGGCCTAGACCAAGACCCAGGCCAAGGCCACGGCCAAGACCTAGA-3’ |
| **101x DPRs** | |
| $GA_{101}$ | 5’GCCGGCGCTGGCGCTGGGGCAGGCGCAGGGGCTGGCGCCGGGGCCGGGGCCGGCGCTGGCGCAGGGGCTGG<br>GGCTGGCGCAGGCGCTGGGGCAGGGGCTGGCGCTGGGGCTGGCGCAGGCGCAGGCGCTGGCGCTGGCGCAGGG<br>GCTGGCGCAGGCGCTGGGGCTGGCGCTGGGGCAGGGGCAGGGGCAGGGGCTGGGGCAGGCGCTGGCGCAGGC<br>GCAGGCGCAGGGGCAGGCGCTGGGGCTGGGGCTGGGGCTGGCGCAGGGGCCGGGGCCGGGGCAGGGGCAGGC<br>GCAGGCGCAGGGGCTGGGGCAGGGGCAGGCGCAGGGGCTGGCGCTGGCGCTGGCGCCGGGGCCGGGGCCGGC<br>GCAGGGGCTGGCGCTGGGGCAGGCGCTGGCGCAGGGGCAGGGGCAGGCGCTGGCGCTGGGGCAGGGGCTGGG<br>GCCGGGGCCGGCGCAGGCGCTGGGGCAGGCGCAGGCGCAGGCGCTGGGGCCGGGGCCGGGGCTGGCGCTGGC<br>GCTGGCGCAGGCGCTGGGGCTGGGGCAGGCGCCGGGGCCGGGGCCGGGGCAGGCGCTGGGGCTGGGGCTGGC<br>GCAGGGGCAGGCGCTGGGGCAGGCGCAGGGGCC-3’ |
| $GR_{101}$ | 5’CGGGGCAGAGGCCGGGGAAGAGGCAGAGGACGCGGAAGGGGAAGGGGGAGAGGAAGAGGGCGGGGACGCG<br>GCCGGGGCCGCGGGCGGGGCCGGGGCCGGGGGCGCGGCCGGGGGCGGGGGCGCGGGCGCGGCCGGGGCCGG<br>GGCCGCGGGCGGGGGCGCGGCCGGGGCCGGGGGCGGGGGCGCGGGCGGGGGCGGGGAAGAGGCAGGGGCAG<br>AGGAAGAGGAAGAGGACGGGGGAGGGGCAGAGGAAGGGGACGCGGCAGAGGCAGGGGACGGGGACGCGGAA<br>GAGGAAGAGGCAGAGGCAGGGGGAGAGGACGGGGCAGGGGGAGGGGCAGGGGACGGGGGAGAGGACGCGG<br>ACGCGGACGCGGCAGGGGAAGAGGAAGGGGAAGGGGAAGAGGACGCGGCAGAGGACGGGGCAGAGGAAGAG<br>GCCGGGGAAGAGGACGCGGAAGGGGGAGAGGCCGCGGAAGAGGAAGGGGAAGAGGGCGGGGACGGGGAAGG<br>GGACGGGGACGGGGCAGAGGCAGAGGAAGGGGCAGGGGCAGGGGAAGGGGCAGAGGAAGAGGAAGGGGAAG<br>AGGCAGAGGAAGAGGACGCGGCAGAGGCCGCGGAAGAGGACGG-3’ |
| $AP_{101}$ | 5’CCCGCCCCTGCCCCTGCTCCAGCTCCTGCACCAGCACCCGCTCCAGCCCCAGCCCCCGCACCCGCCCCTGCACCCG<br>CACCAGCACCAGCCCCTGCTCCAGCCCCCGCTCCTGCTCCTGCCCCAGCCCCAGCTCCCGCTCCAGCTCCTGCTCCCG<br>CTCCAGCCCCTGCACCAGCCCCTGCCCCCGCTCCCGCACCAGCTCCAGCACCAGCTCCCGCCCCTGCTCCAGCACCAG<br>CCCCAGCACCAGCACCAGCTCCCGCACCAGCCCCTGCCCCAGCTCCTGCTCCCGCCCCTGCTCCTGCCCCCGCACCCG<br>CACCCGCTCCCGCCCCAGCTCCAGCTCCTGCCCCTGCCCCCGCTCCAGCACCCGCCCCAGCTCCCGCTCCTGCACCCG<br>CTCCCGCTCCCGCACCCGCCCCAGCCCCTGCACCAGCTCCAGCTCCAGCTCCTGCTCCTGCACCAGCTCCCGCACCCG<br>CCCCAGCTCCAGCTCCCGCTCCAGCACCAGCCCCTGCCCCTGCCCCAGCCCCTGCCCCTGCACCAGCACCAGCACCC<br>GCTCCAGCACCCGCACCAGCCCCAGCACCCGCCCCTGCACCAGCTCCCGCCCCAGCCCCCGCTCCT-3’ |
| $PR_{101}$ | 5’CGGCCCAGACCCCGGCCTAGACCAAGACCCAGACCAAGGCCAAGACCTCGGCCTCGCCCTAGGCCACGCCCTCGC<br>CCCAGACCTAGACCTAGGCCTCGGCCAAGACCTAGGCCCAGGCCTAGACCCCGCCCACGCCCTAGACCACGGCCAA<br>GGCCTAGACCTAGACCAAGGCCCAGGCCCAGACCAAGGCCTCGCCCCAGGCCAAGACCACGGCCAAGACCAAGAC<br>CACGCCCCAGACCCAGACCCAGACCTAGACCACGCCCTAGGCCAAGGCCTCGGCCACGGCCTCGGCCTAGACCCAG<br>GCCAAGACCTAGACCTCGGCCTAGACCACGGCCTCGCCCACGCCCAAGGCCAAGACCAAGACCTAGACCCCGGCCT<br>CGCCCAAGGCCCAGACCTCGCCCTAGACCTCGCCCAAGACCAAGGCCTAGACCTCGCCCACGGCCCAGACCTAGAC<br>CAAGACCACGGCCACGCCCTAGACCTAGGCCCAGACCTCGGCCCAGACCCAGACCACGGCCTAGACCTAGGCCAAG<br>GCCACGCCCAAGGCCCAGGCCAAGGCCCAGACCAAGACCAAGACCCAGACCTCGGCCAAGGCCAAGGCCAAGACC<br>AAGG-3’ |

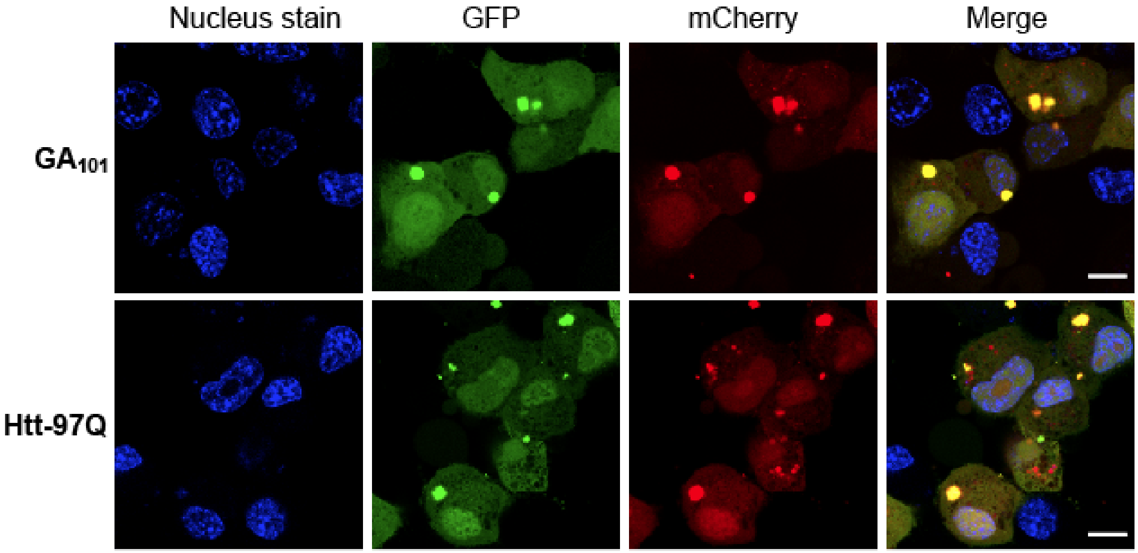


**Fig S1: Expression of DPRs in the ribosome stall reporter constructs. Relates to Fig 3.** Shown are confocal micrographs of Neuro2a cells overexpressing the reporter construct containing either $GA_{101}$ (top panel) or Httex1 with 97Q repeats (bottom panel) fixed 48 hr post-transfection and stained with Hoechst 33258 (blue) to visualize nuclei. Scale bars represent 10 µm.

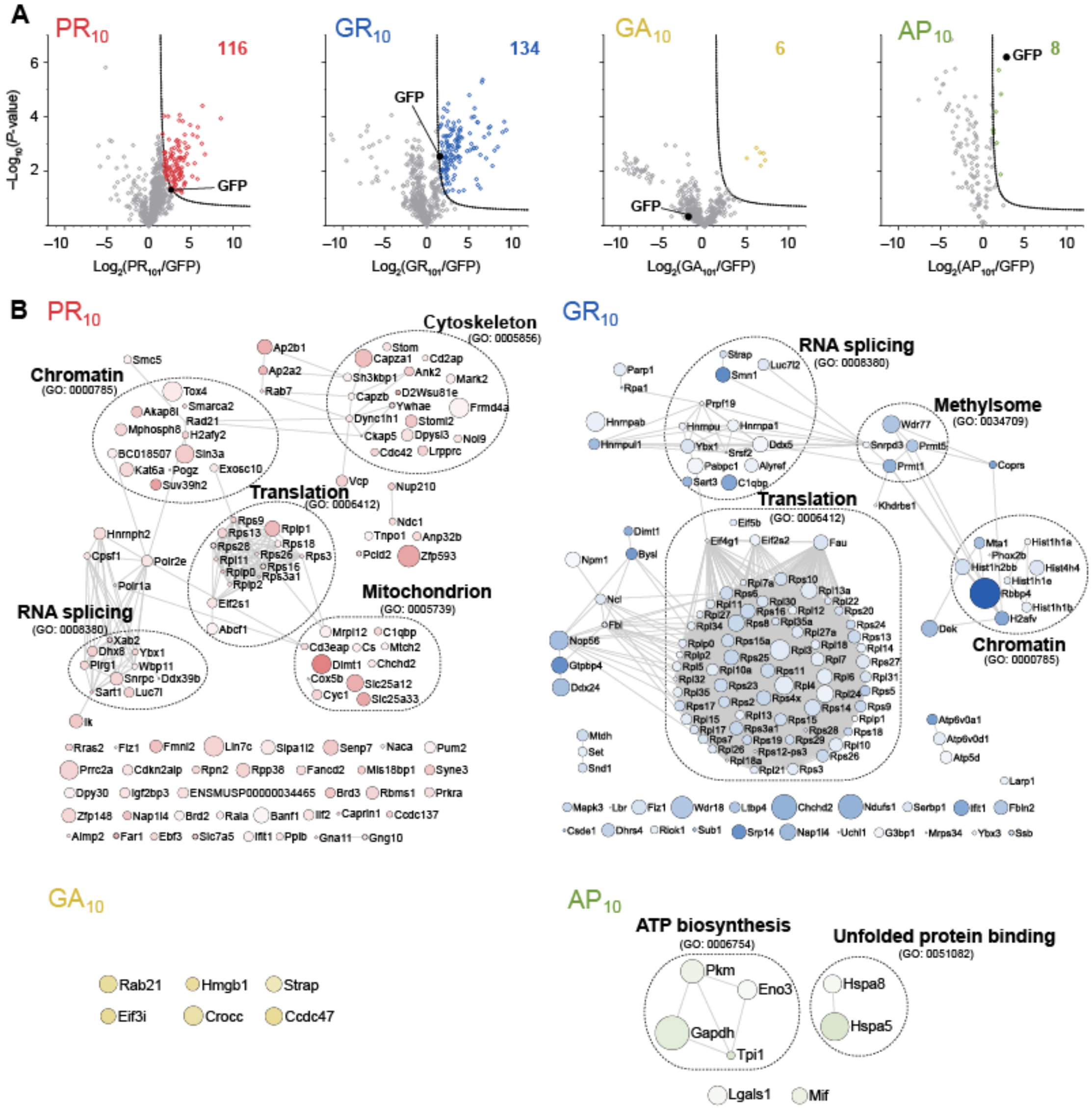

**Fig S2. Interactome analysis of the $DPR_{10}$ variants. Relates to Fig 2. A.** Volcano Plots of each $DPR_{10}$-GFP versus GFP-only control of quantitative proteomics analysis of GFP-Trap immunoprecipitates of $DPR_{10}$-GFP transfected in Neuro2a cells harvested 48h after transfection. Significant binders (shown in colored circles) were classified with False Discovery Rate of ≤ 0.01 (dotted lines). The number of interactors are indicated. **B.** STRING (v10) interaction maps for proteins significantly enriched with confidence set at 0.9 (highest stringency). Circle sizes are proportional to $-\log_{10}$ (*P*-value). The color intensity is proportion to the log (fold change). Selected significantly enriched GO terms (GOCC, GOPB, and UniProt keywords) are displayed (with FDR cutoff *P* value < 0.05).

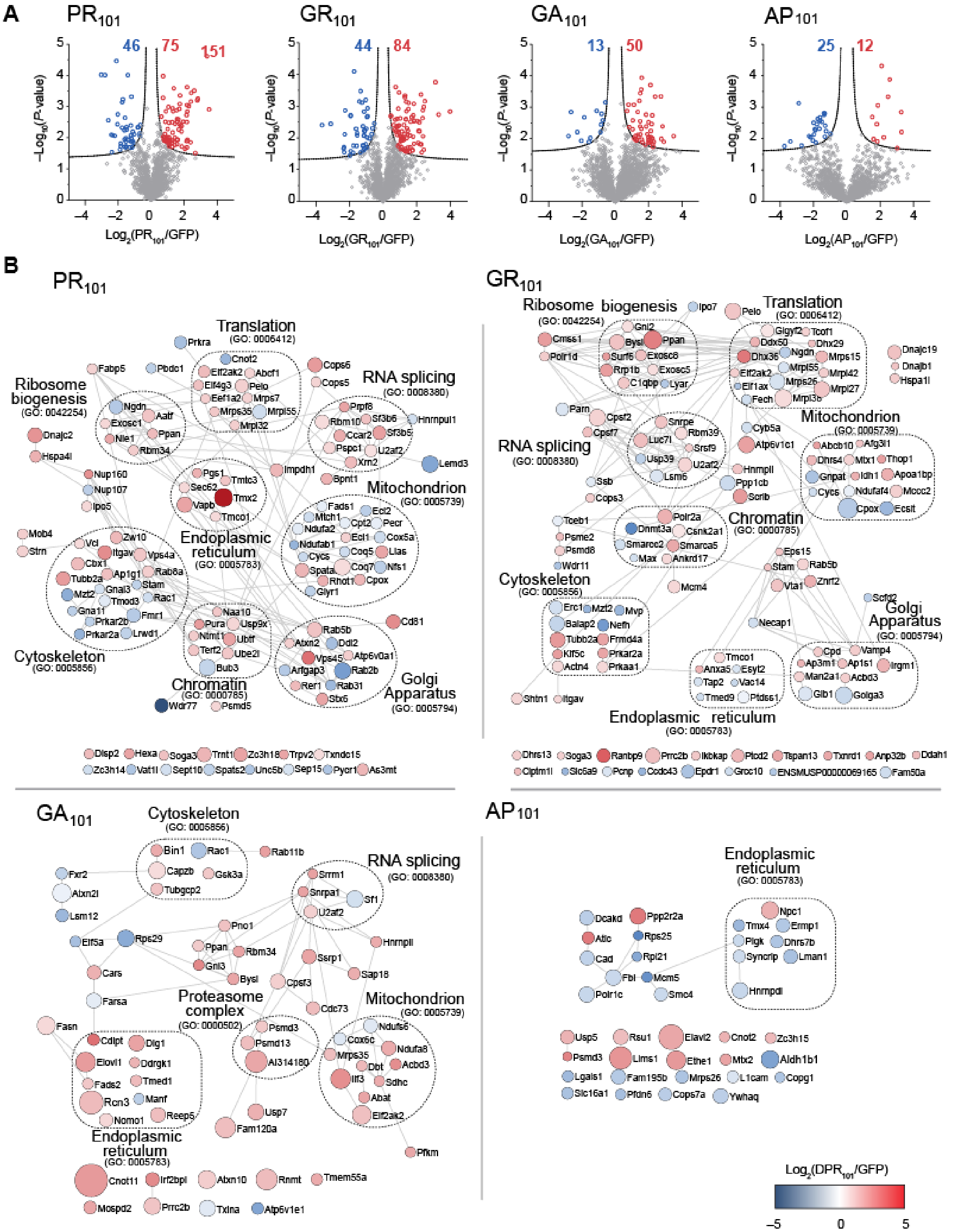


**Fig S3: Proteome abundance changes in response to DPR$_{101}$ expression. Relates to Fig 4.** Shown are protein abundances in Neuro2a cells transfected with each GFP-DPR$_{101}$ compared to GFP for 48 h, and sorted for matched GFP abundance by flow cytometry. *N*=3 biological replicates. *P*-values were determined with a two-sided Welch's t-Test. Significantly changed proteins (red and blue) were defined with FDR ≤ 0.01, $S_0$=1 (dashed lines). The total number of proteins changed are indicated. **B.** Protein interaction networks for proteins significantly changed in abundance, colour coded to enrichment using

STRING (v10) in Cytoscape (v3.6) at medium confidence. Size of nodes scale proportionally to $-\log_{10}$ (*P*-value). Selected significantly enriched GO terms (GOCC, GOPB, and UniProt keywords) are displayed.

**Supplementary References**


1. Abramoff MD, Magalhaes PJ, & Ram SJ (2004) Image Processing with ImageJ. *Biophotonics International* 11(7):36-42.

2. Arrasate M, Mitra S, Schweitzer ES, Segal MR, & Finkbeiner S (2004) Inclusion body formation reduces levels of mutant huntingtin and the risk of neuronal death. *Nature* 431(7010):805-810.

3. Strebel A, Harr T, Bachmann F, Wernli M, & Erb P (2001) Green fluorescent protein as a novel tool to measure apoptosis and necrosis. *Cytometry* 43(2):126-133.

4. Juszkiewicz S & Hegde RS (2017) Initiation of Quality Control during Poly(A) Translation Requires Site-Specific Ribosome Ubiquitination. *Mol. Cell* 65(4):743-750 e744.

5. Grosse R, Copeland JW, Newsome TP, Way M, & Treisman R (2003) A role for VASP in RhoA-Diaphanous signalling to actin dynamics and SRF activity. *EMBO J.* 22(12):3050-3061.

6. Perez-Riverol Y*, et al.* (2019) The PRIDE database and related tools and resources in 2019: improving support for quantification data. *Nucleic Acids Res* 47(D1):D442-d450.
